# Deciphering the non-coding code of pathogenicity and sexual differentiation in the human malaria parasite

**DOI:** 10.1101/2022.10.12.511630

**Authors:** Gayani Batugedara, Xueqing M. Lu, Steven Abel, Zeinab Chahine, Borislav Hristov, Desiree Williams, Thomas Hollin, Tina Wang, Anthony Cort, Todd Lenz, Trevor Thompson, Jacques Prudhomme, Abhai K. Tripathi, Guoyue Xu, Juliana Cudini, Sunil Dogga, Mara Lawniczak, William Stafford Noble, Photini Sinnis, Karine G. Le Roch

**Author notes:** These authors contributed equally to this work (Gayani Batugedara, Xueqing M. Lu, Steven Abel, Zeinab Chahine, Karine G. Le Roch). Corresponding author: Karine G Le Roch, Institute for Integrative Genome Biology, Center for Infectious Disease and Vector Research, Department of Molecular Cell and Systems Biology, University of California Riverside, 900 University Avenue, Riverside CA 92521, USA.

## Abstract

The complex life cycle of *Plasmodium falciparum* requires coordinated gene expression regulation to allow host cell invasion, transmission, and immune evasion. However, this cascade of transcripts is unlikely to be regulated by the limited number of identified parasite-specific transcription factors. Increasing evidence now suggests a major role for epigenetic mechanisms in gene expression in the parasite. In eukaryotes, many lncRNAs have been identified and shown to be pivotal regulators of genome structure and gene expression. To investigate the regulatory roles of lncRNAs in *P. falciparum* we explored the intergenic lncRNA distribution in nuclear and cytoplasmic subcellular locations. Using nascent RNA expression profiles, we identified a total of 1,768 lncRNAs, of which 58% were identified as novel lncRNAs in *P. falciparum*. The subcellular localization and stage-specific expression of several putative lncRNAs were validated using RNA fluorescence in situ hybridization (RNA-FISH). Additionally, the genome-wide occupancy of several candidate nuclear lncRNAs was explored using Chromatin Isolation by RNA Purification (ChIRP). ChIRP-seq of candidate lncRNAs revealed that lncRNA occupancy sites within the parasite genome are focal and sequence-specific with a particular enrichment for several parasite-specific gene families, including those involved in pathogenesis, erythrocyte remodeling, and regulation of sexual differentiation. We further validated the function of one specific lncRNA (lncRNA-ch14) using the CRISPR-Cas9 genome editing tool. Genomic and phenotypic analysis of the △lncRNA-ch14 line demonstrated the importance of this lncRNA in sexual differentiation and sexual reproduction. Our findings bring a new level of insight into the role of lncRNAs in pathogenicity, gene regulation and sexual differentiation. These findings also open new avenues for targeted approaches towards therapeutic strategies against the deadly malaria parasite.

## Introduction

Malaria, a mosquito-borne infectious disease, is caused by protozoan parasites of the genus *Plasmodium*. Among the human-infecting species, *Plasmodium falciparum* is the most prevalent and deadly, with an estimated 627 000 deaths in 2020^1^. The parasite has a complex life cycle involving multiple biological stages in both human and mosquito hosts. This multi-stage developmental cycle is tightly regulated by coordinated changes in gene expression but the exact mechanisms regulating these events are largely unknown.

Compared to other eukaryotes with a similar genome size, *P. falciparum* has an extremely AT-rich genome and a relatively low number of sequence-specific transcription factors (TFs), approximately two-thirds of the TFs expected based on the size of the genome. Our understanding of the regulation of these TFs, and how various TFs could act together to organize transcriptional networks, is still limited. Nascent RNA expression profiles^2^, as well as single cell sequencing^3^ revealed that a majority of the genes in the parasite are transcribed during the trophozoite and schizont stages and that the cascade of gene expression observed using messenger RNA (mRNA) is likely the result of a combination of transcriptional^3-6^ and post-transcription regulatory events^7-10^. Additionally, chromosome conformation capture methods (Hi-C) suggest that the three-dimensional (3D) genome structure of *P. falciparum* throughout its life cycle is strongly connected with transcriptional activity of specific gene families^11^. Therefore, understanding the mechanisms regulating the parasite replication cycle inside the red blood cell (RBC) is essential for identifying key therapeutic targets. Several studies have highlighted the parasite’s ability to escape the host immune response by expressing variants of antigens on the RBC surface^12,13^. To date, several multi-gene families such as *var, rifin* and *stevor*, which encode for *P. falciparum* erythrocyte membrane protein 1 (PfEMP1), RIFIN and STEVOR, respectively, have been identified as key regulators of antigenic variation. In addition, there have been few reports regarding the role of non-protein coding transcripts in the regulation of parasite virulence^14,15^.

With advances in biotechnology and next generation sequencing technologies, huge strides have been made in genomics studies revealing that the transcriptome of an organism is much larger than expected. In eukaryotes spanning from yeast to human, many non-coding RNAs (ncRNAs) have been recently recognized as key regulators of chromatin states and gene expression^16-18^. One class of ncRNAs, the long noncoding RNAs (lncRNAs), are defined as non-protein coding RNA molecules which are ≥ 200 nucleotides in length. Many lncRNAs share features with mature mRNAs including 5’ caps, polyadenylated tails, and introns. In addition, lncRNAs are often expressed and functionally associated in a cell-type specific manner. LncRNAs enriched in the nuclear fraction often associate with regulation of epigenetics and transcription^19-22^, while lncRNAs enriched in the cytoplasm are associated with mRNA processing, post-transcriptional regulation, translational regulation, and cellular signaling process^23-25^. The X inactive specific transcript (Xist), a well-studied example of a lncRNA in mammals that regulates gene expression, mediates X-chromosome inactivation during zygotic development^26^. Deposition of Xist on the X chromosome recruits histone-modifying enzymes that place repressive histone marks, such as H3K9 and H3K27 methylation, leading to gene silencing and the formation of heterochromatin. Similarly, long telomeric repeat-containing lncRNAs (TERRA) have been recently identified as a major component of telomeric heterochromatin^27,28^.

To date, several studies have explored ncRNAs in *P. falciparum* using different techniques^29-31^. Specifically, ncRNAs have been linked to regulation of virulence genes^14,32,33^, and more recently it was established that GC-rich ncRNAs serve as epigenetic regulatory elements that play a role in activating *var* gene transcription as well as several other clonally variant gene families^34^. In addition, a novel family of twenty-two lncRNAs transcribed from the telomere-associated repetitive elements (TAREs) was identified in the parasite^30,33,35^. These TARE-lncRNAs show functional similarities to the eukaryotic family of non-coding RNAs involved in telomere and heterochromatin maintenance^36^. However, chromatin occupancy sites of many lncRNAs in the parasite are still unknown; therefore, the regulatory roles of a large portion of the non-coding transcriptome of *P. falciparum* remain a mystery.

To investigate the regulatory roles of lncRNAs in *P. falciparum* we explored the intergenic lncRNA distribution separately in nuclear and cytoplasmic subcellular locations. Using nascent RNA expression profiles^2^, we identified a total of 1,768 lncRNAs, of which 58% were identified as novel lncRNAs in *P. falciparum*. We further validated the subcellular localization and stage-specific expression of several putative lncRNAs using RNA fluorescence in situ hybridization (RNA-FISH) and single-cell RNA sequencing (scRNA-seq). Additionally, the genome-wide occupancy of several candidate nuclear lncRNAs was explored using Chromatin Isolation by RNA Purification (ChIRP) technology, a method that specifically captures lncRNA-chromatin interactions with high sensitivity and low background. ChIRP-seq of candidate lncRNAs revealed that lncRNA occupancy sites within the parasite genome are sequence-specific with a particular enrichment for several parasite-specific gene families, including those involved in pathogenesis, remodeling of the RBC, and regulation of sexual differentiation. We also demonstrated that the presence of some of these lncRNAs correlates with changes in gene expression. We further validated the role of lncRNA-ch14 using the CRISPR-cas9 editing tool. Functional analysis demonstrated that lncRNA-ch14 is essential during sexual differentiation and development, particularly affecting female gametocytes. Transmission studies demonstrated that even partial deletion of this lncRNA significantly affects parasite development throughout all mosquito stages. Collectively, our results provide crucial information regarding the role of lncRNAs in gene expression and life cycle progression in malaria parasites.

## Results

### Identification of lncRNAs

To comprehensively identify lncRNA populations in *P. falciparum* we extracted total RNA from both nuclear and cytoplasmic fractions using synchronized parasite cultures at early ring, early trophozoite, late schizont, and gametocyte stages (**Fig. 1a**). The samples collected here allow for gene expression profiling during the critical processes of parasite egress, invasion, and sexual differentiation. In brief, extracted parasites were subjected to a modified cell fractionation procedure described in the PARIS kit (ThermoFisher) (see methods). Successful isolation of both subcellular fractions was validated using western blot with an anti-histone H3 antibody as a nuclear marker and an anti-aldolase antibody as a cytoplasmic marker (**Fig. 1b**). After separation of nuclear material from the cytoplasmic material, total RNA and subsequent polyadenylated mRNA was isolated from both fractions. Strand-specific libraries were then prepared and sequenced (see methods for details). For verification, Spearman correlations in gene expression levels were calculated among nuclear samples, cytoplasmic samples, and a previously published steady-state total mRNA dataset generated in our lab^37^ (**Fig. S1**). Once validated, a computational pipeline was implemented for the identification of lncRNAs. Briefly, all nuclear and cytoplasmic RNA libraries were merged, resulting in one nuclear and one cytoplasmic merged file, then assembled into nuclear and cytosol transcriptomes independently using cufflinks. Subsequently, transcripts were filtered based on length, expression level, presence of primary transcript from our previously published GRO-seq dataset^2^, and sequence coding potential (**Fig. 1a**). To specifically identify lncRNA candidates within the intergenic regions, we removed any predicted transcripts that largely overlapped with annotated genes. Our goal was to select transcripts that are ≥ 200 bp in length, consistently expressed in both published nascent RNA and steady-state RNA expression profiles, and that are likely to be non-protein-coding genes. As a result, we identified a total of 1,768 lncRNAs in *P. falciparum* irrespective of the developmental stage. Seven hundred forty-two lncRNAs (42%) overlapped with previously identified intergenic lncRNAs^29,38^, and 1,026 lncRNAs were identified as novel in *P. falciparum* (**Fig. 1c and Supplementary Table S1)**.

**Fig. 1:**
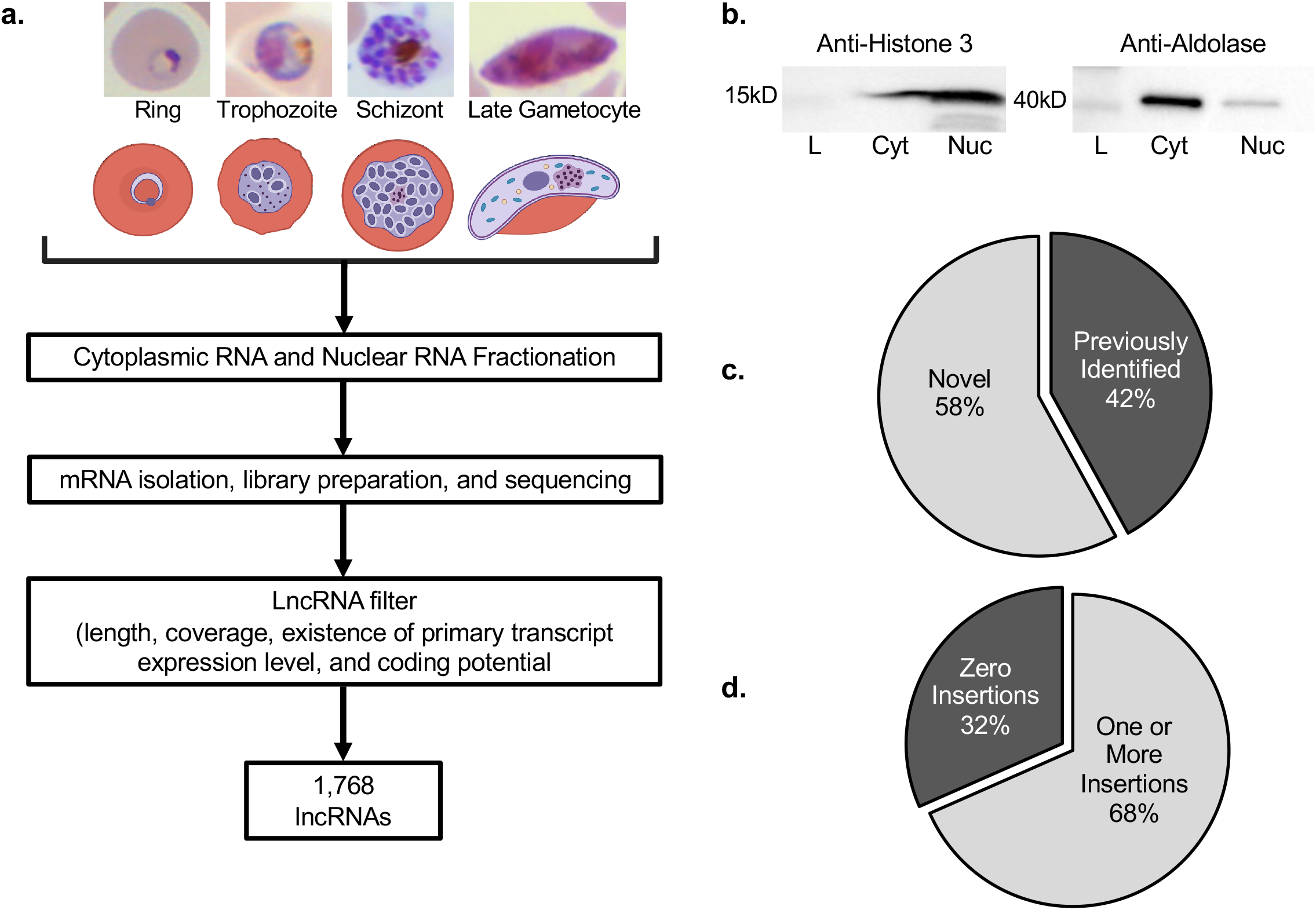
Nuclear and cytoplasmic lncRNA identification. (a) A general overview of the lncRNA identification pipeline. (b) Validation of cell fractionation efficiency using anti-histone H3 and anti-aldolase as nuclear (Nuc) and cytoplasmic (Cyt) markers (L represent the ladder). (c) Comparison of lncRNA candidates with lncRNAs identified from previous publications. (d) Essentiality of lncRNAs using piggyBac insertion (Zhang et al., 2018). LncRNAs that cannot be disrupted are more likely to be essential.

To evaluate the essentiality of the lncRNAs identified in this study, we used piggyBac insertion sites from^39^. In this work, Zhang and colleagues used a high-throughput transposon insertional mutagenesis method to distinguish essential and dispensable genes in the *P. falciparum* genome during the asexual stages of the parasite life cycle. We focused our analysis on the integration of the transposon that occurred within each of the identified lncRNAs. The piggyBac insertion site coordinates were overlapped with the genomic ranges for all detected lncRNAs. This was performed after accounting for differences in the parasite strains used in the two studies. Overall, we were unable to uncover piggyBac insertion for five hundred fifty-eight lncRNAs (31.6%), suggesting that these lncRNAs are potentially essential for the parasite asexual development (**Fig. 1d**). While we observed an insertion in 292 lncRNAs (16.5%) that were specifically detected in gametocyte stage, it will be important to validate at the phenotypic level whether or not those lncRNAs are essential during sexual differentiation. Additionally, significantly fewer insertions per possible insertion site (TTAA sequence) were found for telomeric as compared to subtelomeric lncRNAs, and for subtelomeric as compared to other lncRNAs (**Fig. S2a**). This suggests that piggyBac insertions in telomeric and subtelomeric lncRNAs are most disruptive for parasite survival and that these lncRNAs are more likely to be essential than others. It was also found that the 5’ flanking regions of lncRNAs (**Fig. S2b**) had more insertions per possible site as compared to the rest of the lncRNAs, suggesting that, as a general trend, these regions of the lncRNAs are the most disposable while the gene body and 3’ flanking regions may be more important for their function. Altogether these results indicate that more than 48% of the identified lncRNAs may be essential for parasite survival.

### Length, GC content, and RNA stability of cytoplasmic and nuclear lncRNAs

Next, we categorized our candidate lncRNAs into nuclear lncRNAs, cytoplasmic lncRNAs, or indistinguishable lncRNAs that are equally distributed in both fractions. Among the total identified 1,768 lncRNAs, 719 lncRNAs (41%) were enriched in the nuclear fraction, 204 lncRNAs (11%) were enriched in the cytoplasmic fraction, and 845 lncRNAs (48%) showed similar distribution between both subcellular fractions (**Fig. 2a**). Further, we explored the physical properties of lncRNAs. We observed that lncRNAs are in general shorter in length and less GC-rich as compared to protein-encoding mRNAs (**Fig. 2b and c**). Using total steady-state mRNA expression profiles and nascent RNA expression profiles, we then estimated the expression levels and stability of the lncRNAs. RNA stability was calculated as the ratio between steady-state mRNA expression levels over nascent RNA expression levels. We discovered that, although the overall cell cycle gene expression pattern of the lncRNAs is similar to the expression pattern of coding mRNAs, lncRNAs are less abundant and less stable than coding mRNAs; nuclear IncRNAs are particularly lowly expressed and unstable as compared to the other two groups of IncRNAs (**Fig. 2d**). These observations are consistent with previous lncRNA annotation studies in human breast cancer cells^40^ and noncoding RNA stability studies in mammalian genomes^41^. Our results suggest that the low expression level and the low stability of these lncRNAs may be the reason why they failed to be detected in previous identification attempts. By taking advantage of primary transcripts detected in our GRO-seq dataset, we significantly improved the sensitivity of lncRNA detection, especially for those localized in the nuclear fraction and expressed at a lower level.

**Fig. 2:**
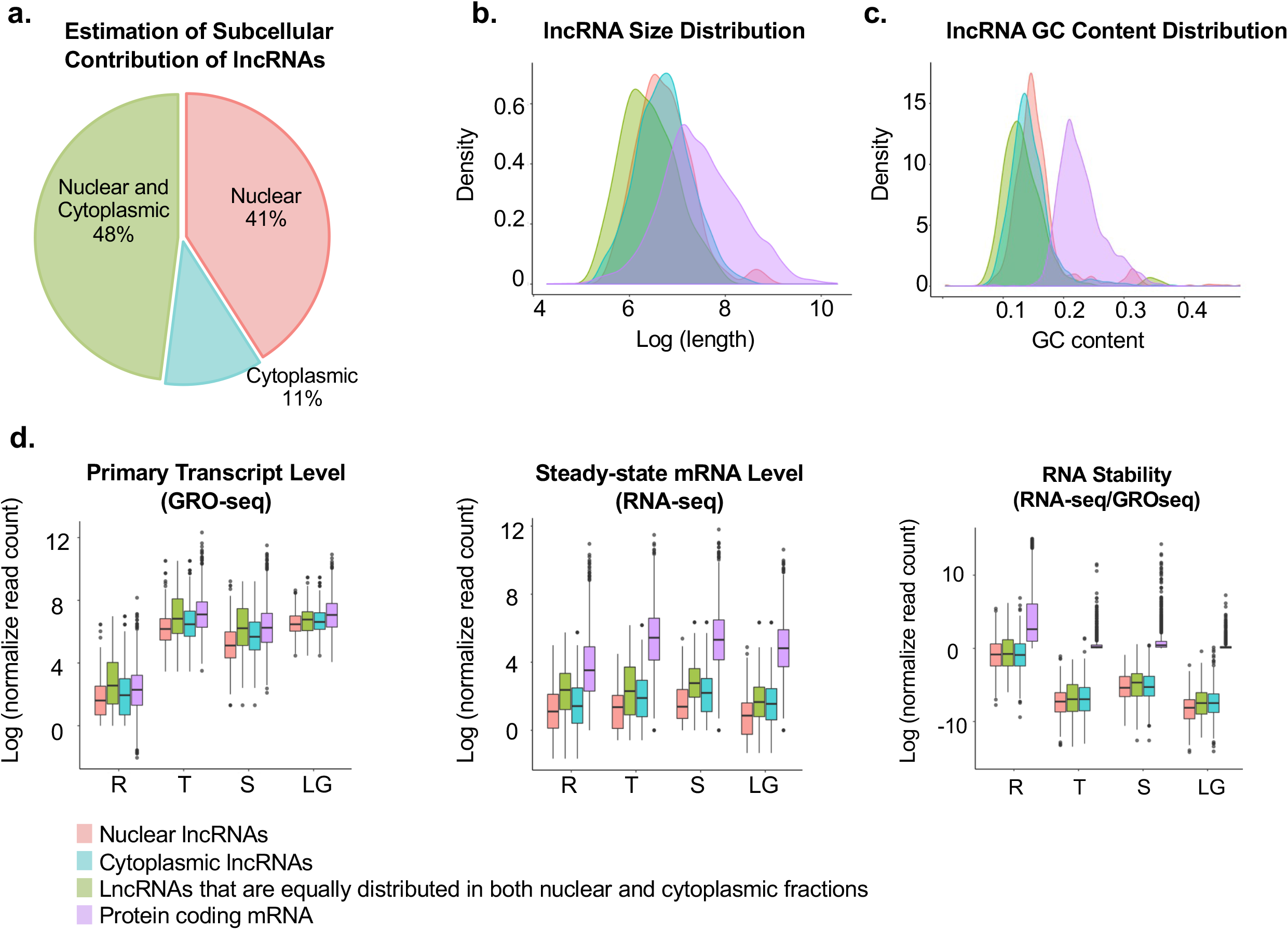
Candidate lncRNA categorization. (a) A total of 1,768 lncRNA candidates were identified, covering 719 nuclear enriched lncRNAs, 204 cytoplasmic enriched lncRNAs, and 845 lncRNAs found in both fractions. Cellular distribution of predicted lncRNAs is based on 0.5 log_2_ fold change of summed nuclear vs cytoplasmic expression level. Density plots of size (b) and GC content (c) of lncRNA candidates and annotated protein encoding mRNAs. (d) Expression levels of primary transcripts (left), steady-state mRNA (middle), and relative stability (right) of lncRNA candidates and annotated protein encoding mRNAs for Ring (R), Trophozoite (T), Schizont (S) and late gametocyte (LG) stages.

### Stage-specific expression of cytosolic and nuclear lncRNAs

As lncRNAs often exhibit specific expression patterns in other eukaryotes, we investigated the stage specificity of identified candidate lncRNAs across the cell cycle. Using k-means clustering, we were able to group lncRNAs into 10 distinct clusters (**Fig. 3a**). Generally, nearly all lncRNAs showed a strong coordinated cascade throughout the parasite’s cell cycle. A large fraction of the lncRNAs was highly expressed at mature stages compared to the ring stages (**Fig. 3b**). Cluster 1 contains lncRNAs that are more abundantly expressed in the nuclear fraction of ring stage parasites and are lowly expressed in the nuclear fraction of schizont stage parasites. LncRNAs representative of this cluster are the lncRNA-TAREs. We observed that a majority of lncRNA-TAREs identified in this study (19 out of 21) are clustered into this group with an average expression of 1.18 log_2_ fold change of nuclear to cytoplasmic ratio (**Fig. 3a**). The remaining two identified lncRNA-TAREs were found in cluster 6, where transcription peaks at the schizont stage. This finding validates our approach and suggests that lncRNAs in this cluster may contribute to the maintenance and regulation of chromatin structure and telomere ends. Approximately 28% of the identified lncRNAs are more abundantly found in either the nuclear or cytoplasmic fraction at the schizont stage (cluster 6, 7 and 8), after DNA replication and the peak of transcriptional activity observed at the trophozoite stage. We observed a few lncRNAs that are solely expressed during the asexual cycle with distinct changes in heterochromatin marks (**Fig. 3c**). The presence of some of these lncRNAs was confirmed using reverse transcriptase PCR (RT-PCR) (**Fig. S3**). Based on clustering analysis, we also found that 25% of the lncRNAs are more exclusively expressed at a high level at the gametocyte stage (cluster 9 and 10). Interestingly, two unique lncRNAs in this cluster, lncRNA-ch9 (Pf3D7_09_v3:1,384,241-1,386,630) and lncRNA-ch14 (Pf3D7_14_v3:3,148,960 - 3,150,115), were identified in a previous study to be located within heterochromatin regions marked by repressive histone marks H3K9me3 at the trophozoite stage^11^ (**Fig. 3d and 3e**). At the gametocyte stage however, the H3K9me3 was lost. Additionally, both lncRNAs are transcribed from regions adjacent to gametocyte-specific genes. To validate the expression of these gametocyte-specific lncRNAs, we performed RT-PCR (**Fig. S3**) as well as single cell RNA-seq (scRNA-seq) across key-stages of the parasite life cycle. LncRNA expression was visualized on the UMAP embedding generated from coding gene expression (**Fig. 3f**). LncRNA-chr9 and lncRNA-chr14 were expressed in sexual-stage parasites, with lncRNA-ch9 and lncRNA-ch14 were expressed in sexual-stage parasites, with specific enrichment in male and female gametocytes, respectively (**Fig. 3f**, right panel). Collectively, these results emphasize the stage-specific expression of parasite lncRNAs and the potential function of gametocyte-specific lncRNAs in regulating sexual development.

**Fig. 3.**
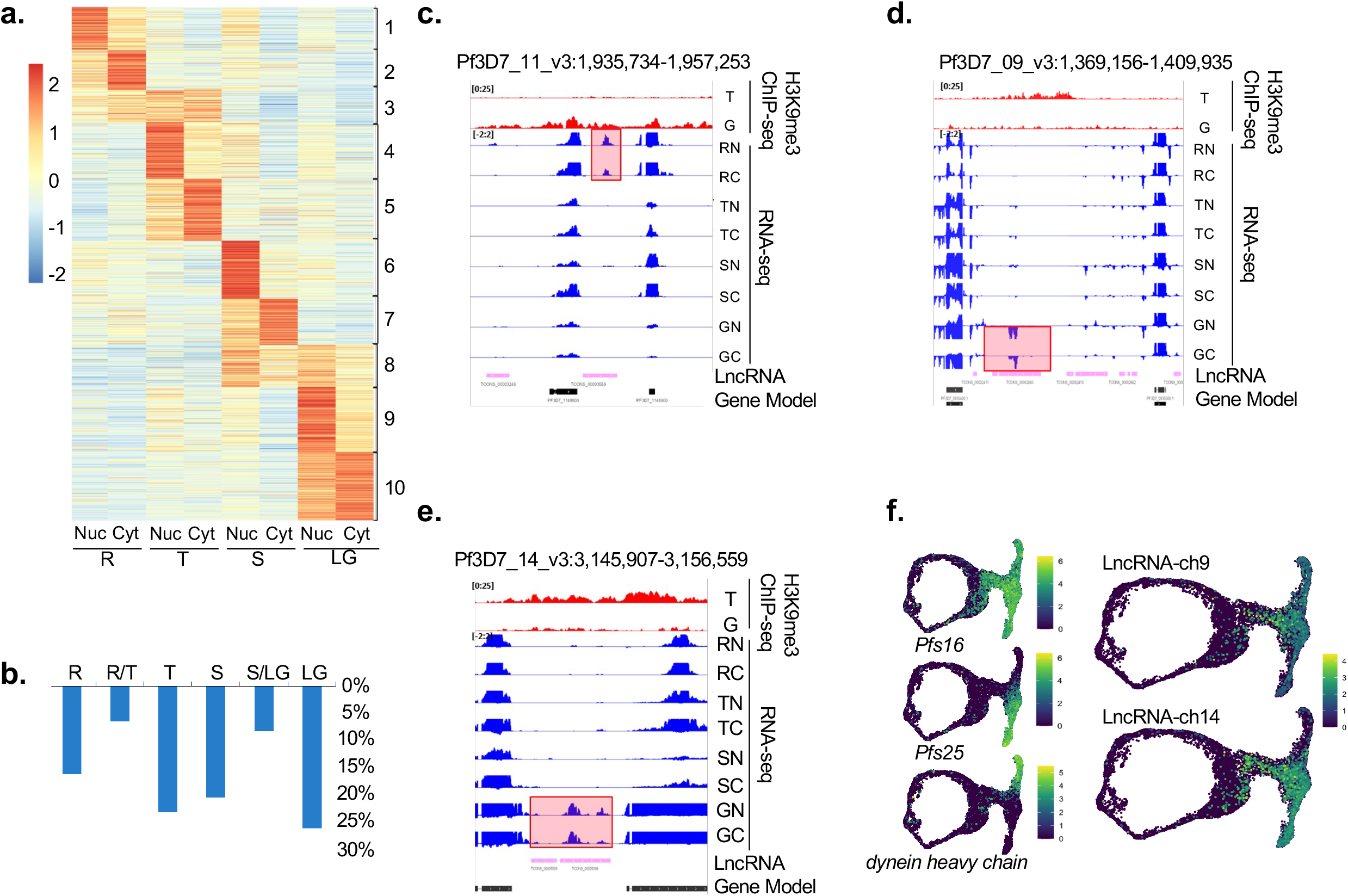
Gene expression pattern of lncRNAs. (a) lncRNAs are grouped into 10 clusters based on their cell cycle expression patterns in the Nuclear (Nuc) or Cytoplasmic (Cyt) fraction for the Ring (R), Trophozoite (T), Schizont (S) and Late Gametocyte (LT) stages (b) Percentage of lncRNAs that are highly expressed at ring, trophozoite, schizont, and late gametocyte stages. Genome browser views of H3K9me3 ChIP-seq and RNA-seq datasets in the region of a representative asexual stage-specific lncRNA on chromosome 11 (c), two gametocyte-specific lncRNAs located at the intergenic regions of chromosome 9, lncRNA-ch9 (d) and 14, lncRNA-ch14 (e). (f) scRNA-seq analysis. 2-dimensional UMAP projection of *P. falciparum* parasites, both asexual (ring) and sexual (T-shape). Each dot represents a single cell. Left panel: Cells colored according to log-normalized gene expression values for gametocytes (*Pfs16* (PF3D7_0406200), top), females (*Pfs25* (PF3D7_1031000), middle), and males (*dynein heavy chain* (PF3D7_0905300), bottom). Right panel: Log-normalized expression of lncRNA-ch9 (top) and lncRNA-ch14 (bottom) across the *P. falciparum* life cycle.

### Validation of lncRNA localization and stage-specific expression

To validate the cellular localization of several candidate lncRNAs, we utilized RNA fluorescence in situ hybridization (RNA-FISH). Two candidate lncRNAs were enriched in the cytoplasmic fraction and 8 lncRNAs were enriched in the nuclear fraction, including one previously identified lncRNA transcribed from the telomere region on chromosome 4, termed lncRNA-TARE4^30^, were used for RNA-FISH (**Fig. 4a and 4b**). Briefly, mixed stage parasites were fixed and hybridized to fluorescently labeled ∼200-300 nucleotide antisense RNA probes (see methods for details). The hybridization images clearly demonstrate that the nuclear lncRNAs localize to distinct foci within the DAPI-stained nuclei (**Fig. 4a**), while cytoplasmic lncRNAs are localized outside the DAPI-stained genomic DNA (**Fig. 4b**). Additionally, using RNA-FISH, we validated the stage-specific expression of our candidate lncRNAs. Specifically, expression of lncRNA-267 (Pf3D7_08_v3: 1382128-1382689) and lncRNA-13 (Pf3D7_01_v3:491225-494291) were enriched at the ring and trophozoite stages; lncRNA-178 was expressed at the trophozoite and schizont stages; lncRNA-643 was expressed at the schizont stage only and lncRNA-TARE4 was expressed at all three asexual stages. LncRNA-ch9 and lncRNA-ch14 were only expressed at the gametocyte stage. These results highlight that, similar to protein-coding transcripts, these candidate lncRNAs are developmentally regulated.

**Fig. 4:**
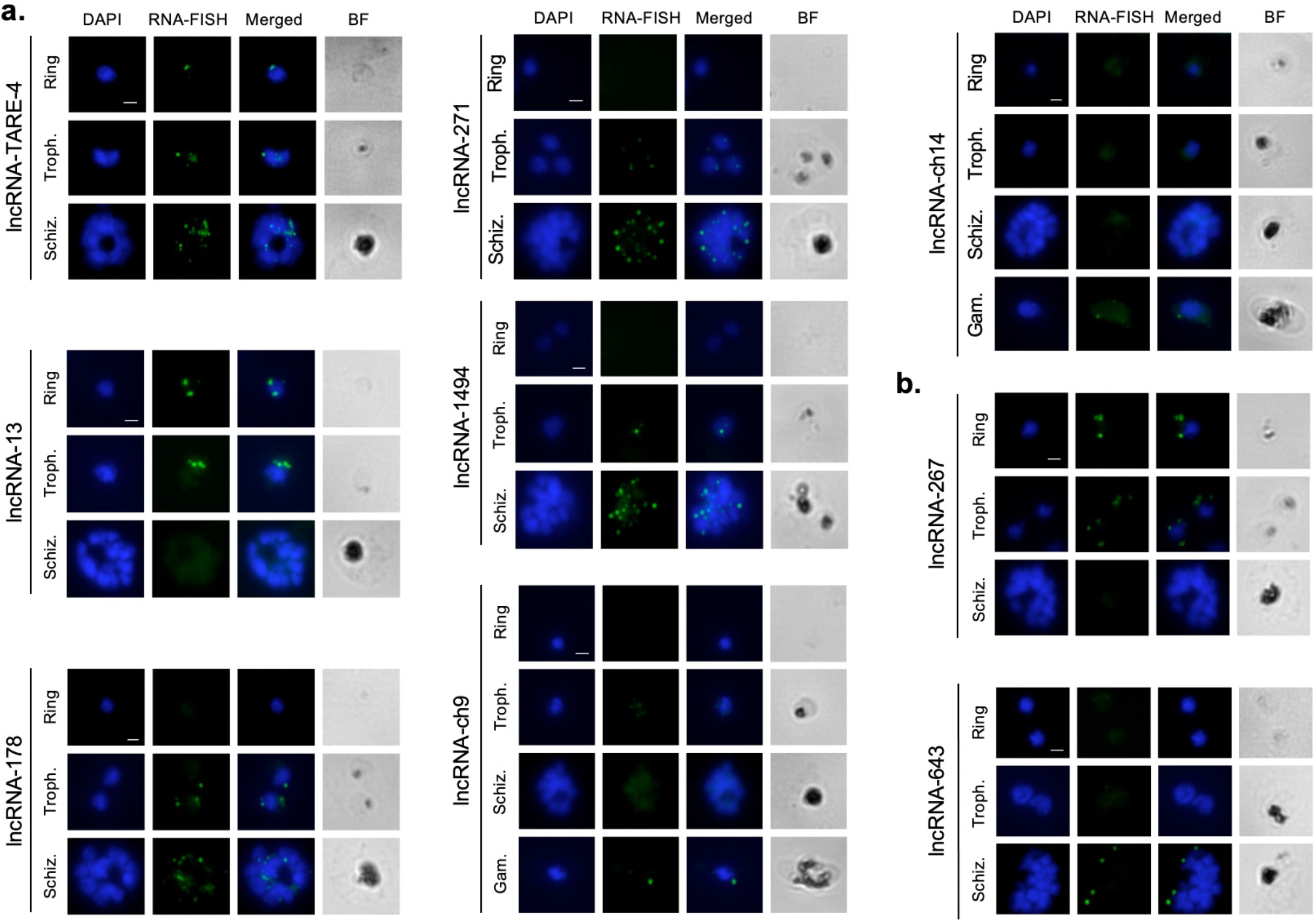
RNA-FISH experiments to show localization of several candidate lncRNAs. Nuclei are stained with DAPI. (a) Nuclear lncRNAs (lncRNA-TARE4, lncRNA-13, lncRNA-178, lncRNA-271, lncRNA-1494, lncRNA-ch9 and lncRNA-ch14) colocalize with DAPI in Ring Trophozoite (Troph.), Schizont (Schiz.) and Gametocyte (Gam.) stages. (b) Cytoplasmic lncRNAs (lncRNA-267 and lncRNA-643) do not colocalize with the nuclei stained with DAPI. Scale bar indicates 2μm.

### Genomic maps of RNA-chromatin interactions

To explore the roles of lncRNAs in chromatin regulation, we sought to identify occupancy sites of our candidate lncRNAs within the parasite genome. For an unbiased high-throughput discovery of RNA-bound DNA in *P. falciparum*, we adapted a method termed Chromatin Isolation by RNA Purification (ChIRP) (**Fig. 5a**)^42,43^. Briefly, synchronized parasites were extracted and crosslinked. Parasite nuclei were then extracted, and chromatin was solubilized and sonicated. Biotinylated antisense oligonucleotides tiling the RNA of interest (**Supplementary Table S1)** were hybridized to target RNAs and isolated using magnetic beads. Purified DNA fragments were sequenced using next-generation sequencing technology. An input control was used to normalize the signal from ChIRP enrichment. Following the pulldown, the RNA fraction was analyzed to validate the specificity of the biotinylated oligonucleotides to target the RNA of interest. RT-PCR results confirmed that lncRNA-TARE4 probes retrieve the lncRNA-TARE-4 and the control serine tRNA ligase probes retrieve the serine tRNA ligase RNA (PF3D7_0717700), respectively (**Fig. 5b**). Neither RNA was retrieved in the negative control that was incubated with no probes. These results confirm that the biotinylated probes target the RNA of interest with specificity.

**Fig. 5:**
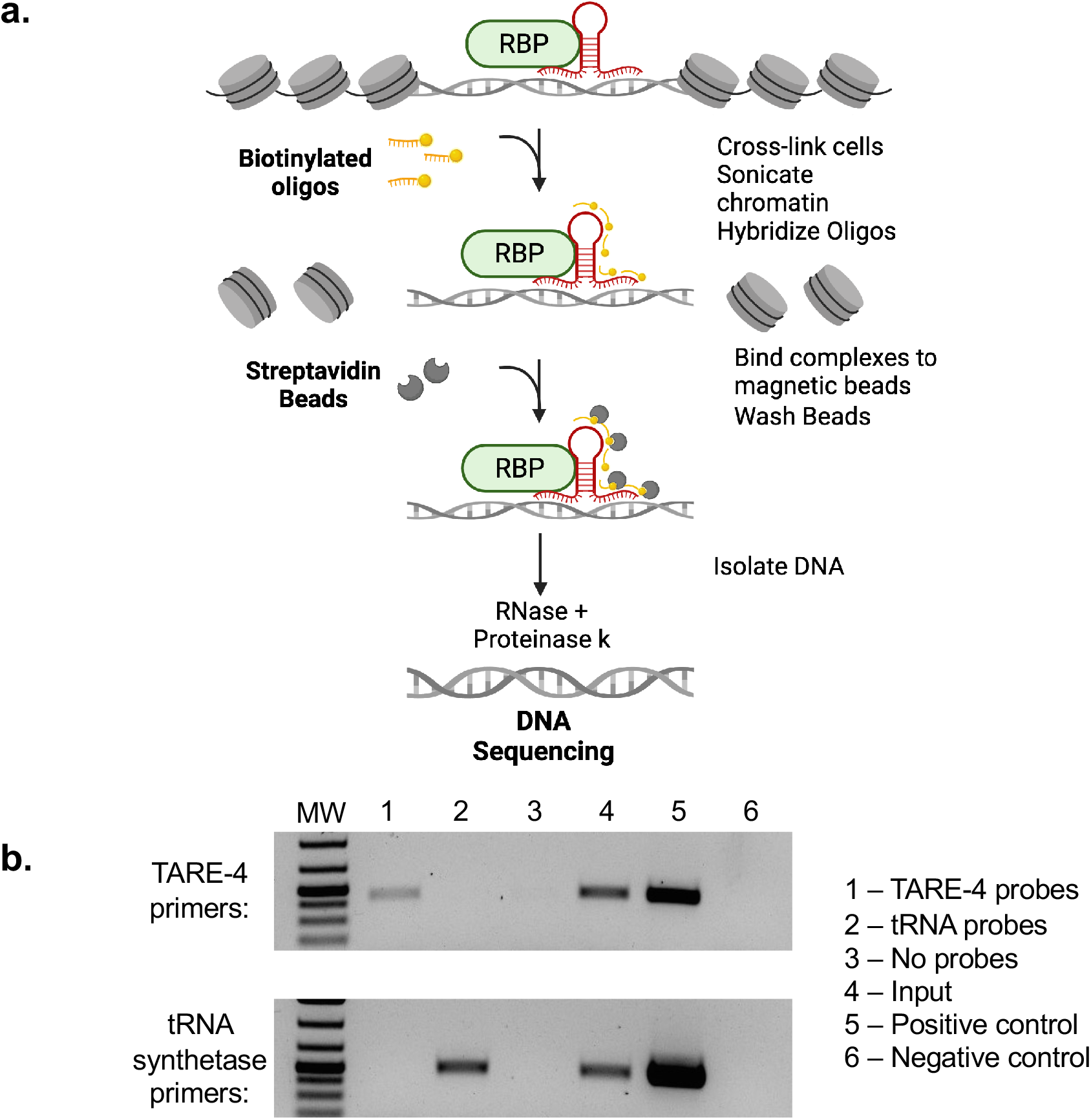
Chromatin Isolation by RNA Purification (ChIRP). (a) Schematic representation of the ChIRP methodology. (b) RT-PCR following the ChIRP protocol validates the specificity of the biotinylated antisense probes. Following the ChIRP-pulldown, the RNA fraction was analyzed, and RT-PCR results confirm that lncRNA-TARE-4 probes retrieve the lncRNA-TARE-4 RNA (535 bp PCR product) and the control serine tRNA ligase probes retrieve the serine tRNA ligase RNA (505 bp PCR product), respectively. Neither RNA was retrieved in the no probe control. RNA from a ChIRP-input sample as well as WT 3D7 parasites (wells 4 and 5, respectively) was used to confirm the lncRNA-TARE-4 and serine tRNA ligase primers. The negative controls (well 6) represent no template controls.

We next wanted to explore the genomic binding sites of seven lncRNAs: lncRNA-TARE4, lncRNA-13, lncRNA-178, lncRNA-1494, lncRNA-271, lncRNA-ch9 and lncRNA-ch14, identified as enriched in the nuclear fraction and validated using RNA-FISH. For all seven lncRNAs, ChIRP-seq experiments were performed in duplicate at the stages the lncRNA was either highly or lowly expressed. For all lncRNAs investigated here, ChIRP-seq performed at time points where the lncRNAs were least expressed retrieved no significant signals (**Fig. 6a, 6b and Fig. S4**). The lncRNA-TARE4, transcribed from the telomere region on chromosome 4, and expressed throughout the parasite life cycle stages investigated in this study was found to strongly interact with most telomeres in a very specific manner **(Fig. 6a**). LncRNA-13, transcribed from a region on chromosome 1 and highly expressed at the trophozoite stage (**Fig. S4**) was enriched around surface antigen genes, including PF3D7_0113100 (SURFIN4.1) and PF3D7_1149200 (RESA, ring-infected erythrocyte surface antigen). Both the SURFIN and RESA families of proteins have been implicated in erythrocyte invasion-related processes and are transcribed in mature-stage parasites^44,45^.Given that a trophozoite stage lncRNA was identified adjacent to surface antigen genes which are transcribed at the schizont stage, lncRNA-13 is possibly playing a role in recruiting chromatin modifying enzymes to edit the epigenetic state of the chromatin and allow the recruitment of transcription factor(s) needed to activate the transcription of these genes. To further investigate the possible link between lncRNAs and their role in epigenetics and regulation of gene expression, we developed a software pipeline in Python to identify all specific binding sites in the genome using Bowtie for mapping and PePr for peak calling^46^. We then aligned lncRNA ChIRP-seq signals across all 5’ and 3’ UTRs as well as the gene bodies. We discovered that the lncRNA occupancy is enriched either in the gene bodies for lncRNA-13 and lncRNA-178 or near the end of the 5′ UTR of each gene (lncRNA-1494 and lncRNA-271) (**Fig. S5**). This pattern provides support for our candidate lncRNAs to promote either transcriptional elongation or transcriptional initiation, respectively. We next retrieved the genes closest to the identified lncRNA binding sites and calculated the log_2_ fold change of their expression from inactive to active stage. We compared the resultant information to the change in the expression profiles for all other genes in the *P. falciparum* genome (**Fig. 6c**). For each of the lncRNAs investigated, we detected a significant increase in the expression of the genes near the ChIRP signals. These results indicate that overall, the presence of lncRNA correlates with a significantly increased gene expression. To further demonstrate that lncRNAs interact specifically with DNA, we looked for motif enrichment (**Fig. 6d**). Motif analysis of ChIRP-seq data revealed one motif for lncRNA-13 (pval=1.8e-3) occurring in 131 of the 138 retrieved lncRNA-13 sequences and two motifs for lncRNA-TARE-4 (pval=3.7e-7 and pval=5.1e-5) occurring in 72% and 61% of the TARE-4 binding sites. These data demonstrate that our ChIRP-seq experiments were highly sensitive and specific, and that we were able to retrieve biological insights into their function.

**Fig. 6:**
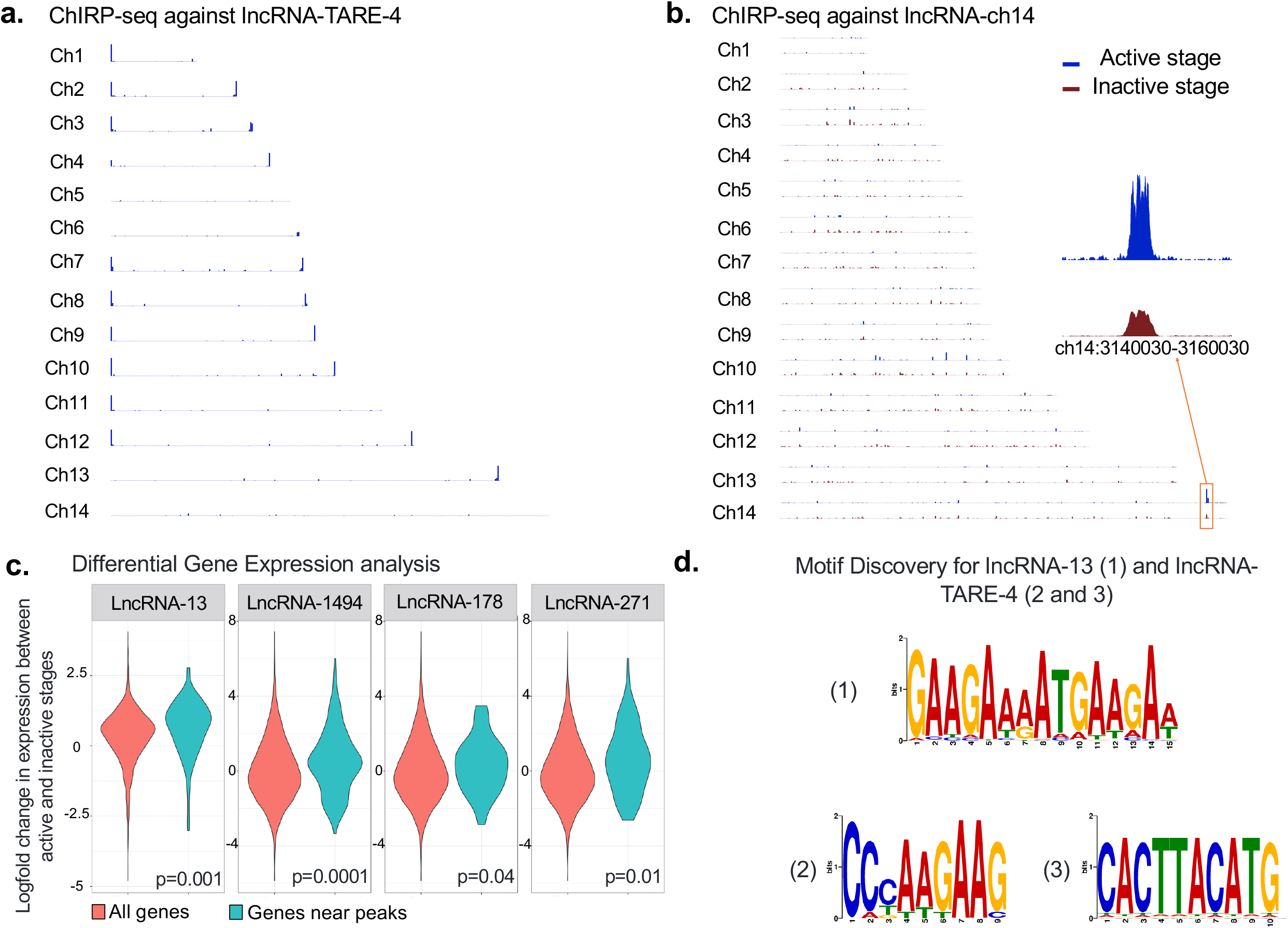
ChIRP-seq reveals candidate lncRNA binding sites. (a) Genome-wide binding sites of lncRNA-TARE-4 and (b) lncRNA-ch14. Mapped, normalized reads from the active stage (top, blue track) and inactive stage (bottom, red track) are shown for each chromosome. (c) Differential gene expression analysis. The log2-fold change of gene expression was calculated for the genes closest to the lncRNA peaks between the inactive and active stage (right violin). This distribution was compared to the log2-fold change in expression for all other genes in the *Plasmodium* genome (left violin) using a t-test, and the p-values are reported at the bottom of each panel. (d) Motif identification. 100bp sequences centered at the peaks’ summits were extracted, and we used STREME specifying 2nd order Markov model and default for the rest of parameters to search for possible motifs. We identified one motif for lncRNA-13 (p=1.8e-3) occurring in 131 of the 138 (95%) lncRNA-13 sequences and two motifs for lncRNA-TARE-4, (p=3.7e-6 and p=5.1e-5), occurring in 72% and 61%, respectively, of the TARE binding areas.

We then focused our attention on ChIRP-seq data generated using probes against lncRNA-ch9 and lncRNA-ch14, two gametocyte-specific lncRNAs. ChIRP-seq signals (**Fig. 6b and Fig. S4**) showed significant enrichment in the genomic regions where the lncRNAs are transcribed. The lncRNA-ch9 lie between genes that have been implicated in gametocyte differentiation. These include PF3D7_0935500^47^, a *Plasmodium* exported protein of unknown function, PF3D7_0935600, a gametocytogenesis-implicated protein and PF3D7_0935400, Gametocyte development protein 1. These three genes are known to be significantly up regulated in gametocytes and have been demonstrated to be essential to sexual commitment^48^. For lncRNA-ch14, the genes are PF3D7_1476500, a probable protein of unknown function, PF3D7_1476600^49^, a *Plasmodium* exported protein of unknown function and PF3D7_1476700, a lysophospholipase, three genes on chromosome 14 that are only detected either in gametocyte or ookinete stages. When overlaid with previous ChIP-seq data generated against histone H3K9me3 during the asexual and sexual stages of the parasite life cycle, we noticed that the presence of these lncRNAs correlate with a loss of H3K9me3 marks at the gametocyte stage. While these data will need to be further validated, our results suggest that these lncRNAs may recruit histone demethylase and/or histone acetyl transferase to change the epigenetic state of the chromatin and activate the expression of these genes during sexual differentiation. Collectively, these experiments propose that lncRNAs in the parasite could be essential to recruit chromatin remodeling and modifying enzymes as well as sequence-specific transcription factors to regulate gene expression.

### Role of lncRNA-ch14 in sexual commitment and development

We next sought to validate the role of one lncRNA in the parasite development. We therefore selected lncRNA-ch14, which was detected as upregulated in female gametocytes. We began by disrupting its full length via the CRISPR-cas9 editing tool. After four unsuccessful attempts, we concluded that disruption of this large genomic region was possibly lethal to the parasite. However, we were able to successfully disrupt the lncRNA-ch14 gene through insertion of a resistance marker spanning the position (Ch14:3,148,960 - 3,150,115) of the gene **(Fig. S6a)**. Parasite lncRNA14 disruptive lines (two clones, named △lncRNA-ch14 B1 and F2) were recovered and validated via PCR and RT-PCR **(Fig. S6b)**. We then examined parasite growth using our two selected clones along with wild-type (WT) NF54 parasites during the erythrocytic cycle. Growth was monitored in triplicates using Giemsa-stained blood culture smears for two full cycles. No significant difference was observed in the asexual stages of the △lncRNA-ch14 clones compared to the WT (data not shown). This indicates that lncRNA-ch14 does not have a significant role in the asexual blood stage replication.

We subsequently aimed to analyze the effect of △lncRNA-ch14 in gametogenesis. Relative gametocyte numbers were determined by microscopic examination of Giemsa-stained blood prepared from day 16 gametocyte culture smears. To consider the impact prolonged culturing times may have on gametogenesis, the assays were conducted between our two △lncRNA-ch14 clones as well as two NF54 strains; a NF54 WT lab strain as well as the NF54 parental line used for the initial transfection. The NF54 parental line was maintained in culture in parallel with our selected △lncRNA-ch14 clones. We detected a significant decrease in mature stage V gametocytes in clones B1 and F2 compared to the WT NF54 line (n=4, p<0.05). This decrease was however not detected as significant between the parental NF54 line and △lncRNA-ch14 clones **(Fig. 7a)**. To better understand discrepancies observed between the NF54 lines, we purified gDNA for whole genome sequencing (WGS). While we confirmed successful disruption of ncRNA-ch14 in our two selected clones, we also identified a nonsense mutation in the gametocyte developmental protein 1 (*gdv1)* gene (PF3D7_0935400), a gene essential in sexual differentiation, in the parental line. The mutation detected indicated a premature stop codon leading to a C-terminal truncation of 39 amino acids (GDV39) similar to what was previously observed by Tibúrcio and colleagues^50,51^. Of the 110 reads covering the mutation site, 43 were shown to retain the reference base while 67 displayed the *gdv1* mutation described. This result suggests the development of a spontaneous mutation after transfection and that the reduced number of gametocytes observed in the parental line were most likely a result of a significant portion (60%) of parasites with the *gdv1* mutation, not producing mature gametocytes. This mutation was however absent in both of our △lncRNA-ch14 clones as well as our NF54 WT **(Supplementary Table S2)**, explaining the discrepancies observed in our gametocyte induction assays between the NF54 parental line. Development of spontaneous mutations in culture attests the need of WGS to validate the phenotypes observed in the parasites for both the WT and genetically modified strains. As the parental line was still capable of generating healthy mature gametocytes, albeit at a lower frequency, and gametocytemia is normalized prior to mosquito feeds, we decided to use the NF54 parental line as our control to analyze the impacts of lncRNA-ch14 disruption on transmission to the mosquito. This decision was made since the gametocytes produced from the parental line, albeit reduced, would more closely resemble the △lncRNA-ch14 clones at the genomic level. Thus, any resulting phenotypic differences observed could be considered more significant as opposed to using an unrelated WT NF54 line.

**Fig. 7:**
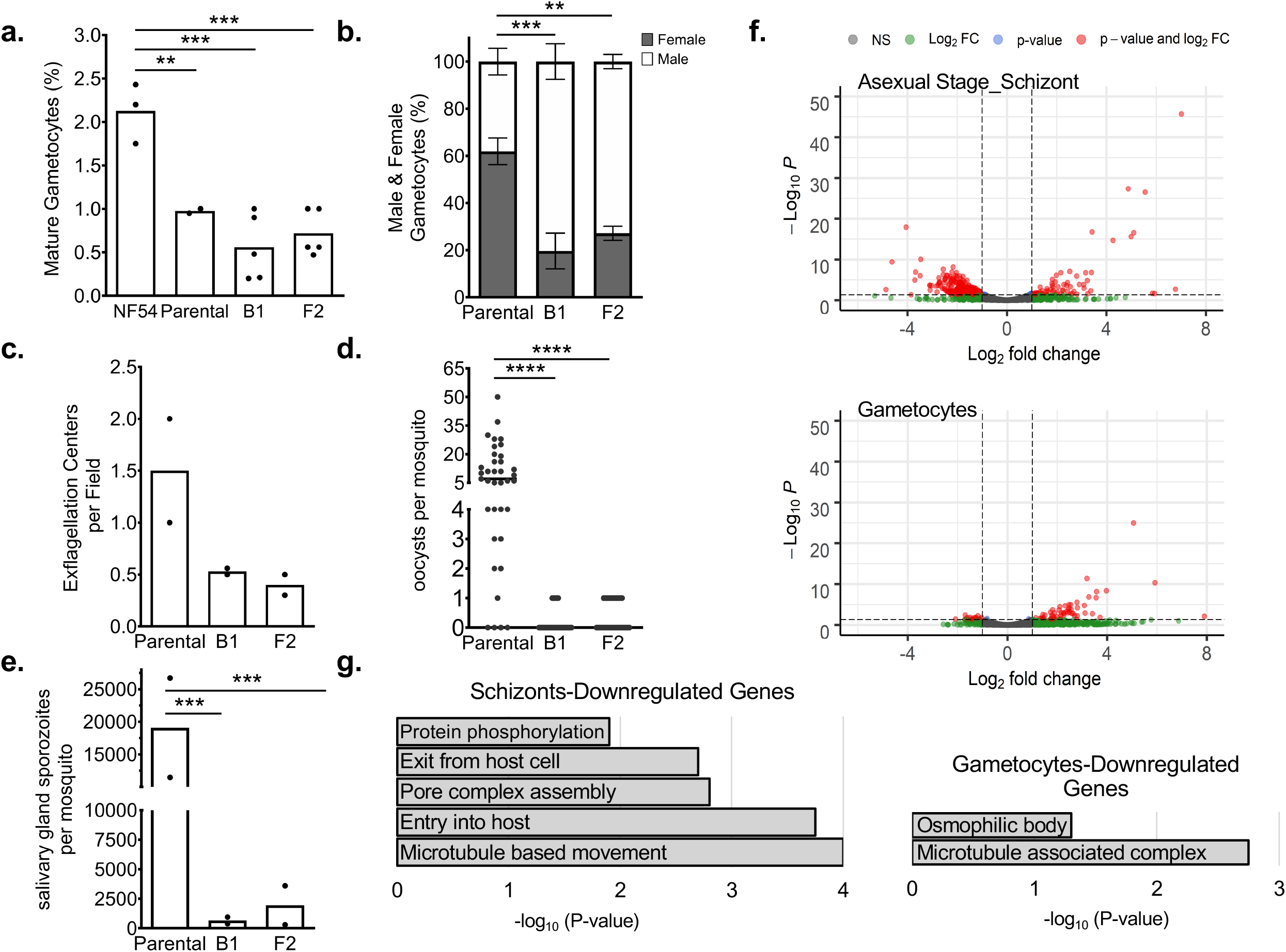
LncRNA-ch14 disruption design and characterization. (a-c) Gametocyte characterization: Gametocyte cultures were sampled by Giemsa-stained blood smears to assess the percent of Stage V gametocytes (a) and the percentage of mature gametocytes identified as male and female (b). Significance of the results were calculated using the One-way ANOVA with Holm-Šídák correction (**p < 0.01, ***p < 0.001) (a) and the Two-way ANOVA and Tukey’s multiple comparison test (**p < 0.01, ***p < 0.001) (b). Exflagellation assays were performed and the number of exflagellation centers per field were counted (c). Shown are results from two biological replicates. Bars indicate the mean. In panel b shown are means +/-SD. (d&e) Mosquito passage: Gametocyte cultures were fed to *Anopheles stephensi* mosquitoes. On day 11 post-infection, midguts were removed, and oocysts were counted (d) and on day 17 post-infection salivary glands were harvested and sporozoites were counted (e). Data are pooled from 2 biological replicates. Oocyst counts were performed with 15 to 25 mosquito midguts per experiment. Significance of the results was calculated using the Kruskal-Wallis test with Dunn’s post-test (****p < 0.0001). For salivary gland sporozoite counts, salivary glands from 20 mosquitoes were harvested, homogenized and sporozoites counted using a hemocytometer. Shown is the average number of salivary gland sporozoites for each of two experiments. Significance was calculated using the Chi-Squared tests (***p <0.001). (f) Volcano plots for gene expression profile between the WT and △lncRNA-ch14 lines by –Log10 P (y-axis) and log_2_ Fold (x-axis) fold change in asexual stage parasites (Top) and in mature gametocytes (Bottom). G. Select gene expression profiles between WT and △lncRNA-ch14 lines are presented in bar graph by Log10P (y-axis) in asexual stage mature parasites (Left) and mature gametocytes (Right).

We then analyzed the formation and ratio of mature male and female gametocytes between the parental and △lncRNA-ch14 clones. To discriminate between male and female gametocytes, blood smears prepared from day 16 gametocyte cultures were stained with Giemsa and ≥100 mature stage V gametocytes were counted to determine sex ratio in each line. Male gametocytes can be distinguished from their female counterparts as they are less elongated with rounder ends and their cytoplasm is distinctly pink while the cytoplasm of female gametocytes, with their large stores of RNA, are dark blue. As shown in **Fig. 7b**, the male to female ratio was significantly affected in the △lncRNA-ch14 clones compared to control parasites. In control lines the ratio of female to male gametocytes was approximately 2:1, with the expected larger number of females than males. In contrast, in the lncRNA-ch14 clones it was approximately 2:7, with significantly more males than females (n=2, p<0.05) **(Fig. 7b)**. These data suggest that lncRNA-ch14 has a role in the ratio of male and female gametocytes produced under our culture conditions. Exflagellation assays revealed a dramatic drop in microgametocyte exflagellation with an average of a 65% decrease in exflagellation centers observed in the △lncRNA-ch14 clones compared to the parental lines, indicating a defect in male gametogenesis and microgamete formation in △lncRNA-ch14 parasites **(Fig. 7c)**. All together these data demonstrate a role of lncRNA-ch14 in gametogenesis.

We next investigated the transmissibility of △lncRNA-ch14 gametocytes to mosquitoes. Infectious blood meals were prepared with parental and △lncRNA-ch14 clones stage V gametocytes. Mosquitoes were fed with 0.2% mature stage V gametocyte infected blood using membrane feeders as described earlier^52^. Mosquito midguts were dissected and analyzed on day 11 post blood feeding and showed a significant decrease in the number of oocysts per midgut in the △lncRNA-ch14 clones compared to the parental control. Both prevalence and intensity of infection were impacted in the △lncRNA-ch14 clones. While 90% of control infected mosquitoes had oocysts, only 17% and 37% of the △lncRNA-ch14 clones, B1 and F2, respectively were positive for oocysts. Additionally, the mosquitoes infected with the △lncRNA-ch14 clones had only 1 oocyst while the control had a median of 7 oocysts (n=2, p<0.05) (**Fig. 7d**). As expected, these lower oocyst numbers resulted in significantly lower salivary gland sporozoite numbers. On day 17 salivary glands were dissected from control and LlncRNA-ch14 infected mosquitoes and the average of number of salivary glands sporozoites from 2 biological replicates was 19,000 for the control line and 667 and 1993 for the LlncRNA-ch14 clones B2 and F1, respectively, a decrease of about 96% (n=2, p<0.05) **(Fig. 7e)**. Altogether, though we could only partially disrupt our candidate lncRNA, we clearly demonstrate that lncRNA-ch14 has an important role in gametocyte development and in the infectivity of these gametocytes for mosquitoes.

### Transcriptome perturbation in asexual and sexual stages

Based on our phenotypic assays, we predicted that perturbation of lncRNA-ch14 would affect parasite gene expression. We therefore performed RNA-seq analysis for two biological replicates each of our WT NF54 and LlncRNA-ch14 clones. For each parasite line, RNA was extracted at both schizont stage before sexual commitment and at the late gametocyte stages after sexual commitment. We observed up-regulation of 78 and 57 genes respectively in the LlncRNA-ch14 lines compared to controls as well as downregulation of 383 and 18 genes respectively in the LlncRNA-ch14 lines compared to controls **(Fig. 7f)**. Gene Ontology (GO) enrichment analysis revealed that up-regulated genes were involved in translational regulation including ribosomes and ribosomal subunits (Pval= 10e-20) that are known to be expressed in a stage specific manner between the human and vector hosts. Downregulated genes were mostly involved in microtubule movement, cell signaling and oxidation-reduction processes that are known to be critical during sexual differentiation **(Fig. 7g, Fig. S7)**. Five specific genes known to be upregulated in female gametocytes were detected to be significantly downregulated in our LlncRNA-ch14 clones compared to our control line. These genes include PF3D7_1250100, the osmiophilic body protein G377; PF3D7_1407000, LCCL domain-containing protein); PF3D7_0719200, the NIMA related kinase 4; PF3D7_0525900, the NIMA related kinase 2, and PF3D7_1031000, the ookinete surface protein P25. Four male-specific genes were also detected as downregulated in the lncRNA-ch14 clones compared to the control. Those genes included PF3D7_1113900, the mitogen-activated protein kinase 2, PF3D7_1014200, the male gamete fusion factor HAP2; PF3D7_1216700, the perforin-like protein 2 and PF3D7_1465800, the putative dynein beta chain coding gene. All together this data confirms that lncRNA-ch14 controls, at least partially, the regulation of the transcripts known to be critical in gametocyte development including several key kinases involved in cell signaling relevant to gametogenesis.

## Discussion

Many lncRNAs are now recognized as essential regulators of chromatin structure and gene expression in eukaryotes. Specifically, functions of nuclear lncRNAs have been determined as either directly promoting or repressing gene expression activity^53,54^, guiding or enhancing the functions of regulatory proteins^20,54-57^, or assisting the alteration of chromatin structures by shaping 3D genome organization ^21,58,59^. Some of the well-characterized nuclear lncRNAs, such as Xist^60^, Firre^61^, and Neat^62^, were shown to be particularly important for nuclear organization and chromatin conformation changes. In the last decade, many lncRNAs have been also discovered with diverse cellular functions outside of the nucleus. These types of lncRNAs have been reported to interact with ribosomes^24^ and are often associated with post-transcriptional and translational processes^23^. Some cytoplasmic lncRNAs, such as half-STAU1-binding site RNAs (1/2-sbsRNAs)^63,64^ and growth arrested DNA-damage inducible gene 7 (gadd7)^65^, are shown to alter the stability of mRNA, while others, including lncRNA-p21^66^ and AS UCHl1^67^ are shown to either promote or repress translational processes.

The extent of lncRNA regulation in the malaria parasite is only now starting to emerge. In *Plasmodium*, a variety of lncRNAs transcribed from telomere-associated repetitive elements (TAREs) have been identified^30^ with suggested roles in telomere replication and structural maintenance. In addition, two lncRNAs transcribed from a bidirectional promoter within the intron of *var* genes have been suggested to function in the regulation of *var* genes^14^. More recently, a family of ncRNAs transcribed in a clonally variant manner was linked to transcriptional regulation of various multigene families involved in parasite virulence^34,68^. A lncRNA, *gdv1* antisense RNA, was also identified as a negative regulator of gdv1 transcription, involved in sexual commitment^48^. Collectively, these studies contribute greatly to our knowledge of previously unidentified *P. falciparum* lncRNA candidates.

The dataset generated in this study presents the first global detection of lncRNAs from different subcellular locations throughout the cell cycle of *P. falciparum*. Using both experimental and computational pipelines, we identified 1,768 lncRNAs covering 204 cytoplasmic enriched, 719 nuclear enriched, and 845 lncRNAs that localized to both fractions. Our data suggest that cytoplasmic lncRNAs are coordinately expressed but are less abundant as compared to the number of nuclear lncRNAs in the parasite. In addition, we observed that a small group of cytoplasmic lncRNAs is highly expressed at the trophozoite stage, the stage where a large proportion of genes are transcribed ^2^. Though more in-depth studies will be required to confirm the functions of these trophozoite-expressed cytoplasmic lncRNAs, it is possible that some of these lncRNAs are involved in mRNA stability, alternative splicing, or translational regulation.

By utilizing published nascent RNA expression profiles (GRO-seq ^2^), we were able to significantly improve the sensitivity of lncRNA detection, especially for the identification of nuclear lncRNAs. In *P. falciparum*, emerging evidence has shown that chromatin structure and genome organization are of vital importance for the parasite’s gene expression and regulation system^11,69^. Therefore, identification and characterization of nuclear enriched lncRNAs may help us to uncover chromatin-associated regulators in the parasite.

In our present work, we observed that a large number of lncRNAs, including the lncRNA-TAREs, are highly abundant at the ring and schizont stages. This finding suggests that some of these lncRNAs (cluster 1, **Fig. 3a**) are likely to be involved in heterochromatin maintenance or chromatin structure re-organization events, as previous Hi-C experiments show that chromatin structure is compacted at the ring and schizont stages^70^. Additionally, ChIRP experiments mapping genome-wide binding sites of lncRNAs revealed that lncRNA-TARE4 binds to subtelomeric regions on multiple chromosomes as well as regulatory regions around genes involved in pathogenesis and immune evasion. Previous reports showed that subtelomeric regions as well as virulence gene families cluster in perinuclear heterochromatin. Therefore, evidence suggests a role for lncRNA-TARE4 in transcriptional and/or epigenetic regulation of parasite telomeric and subtelomeric regions by interacting with or recruiting histone-modifying complexes to targeted regions in order to maintain them in a heterochromatin state, much like the case of X chromosome inactivation regulated via lncRNA Xist^60^.

Genomic occupancy of other lncRNAs explored here, suggest that these lncRNAs bind around the gene regions (**Fig. 6**). In all cases investigated, a positive correlation was observed between the lncRNA expression and the expression of genes around the lncRNA occupancy sites. Given already existing evidence for lncRNA-associated epigenetic modification and transcriptional regulation in other eukaryotes^71-73^, it is likely that the lncRNAs identified in the parasite nucleus are responsible for coordinated recruitment of distinct repressing proteins and/or histone-modifying complexes to target loci. Additionally, we uncovered that the lncRNA binding sites were situated upstream of the start codon of target genes (**Fig. S5**). This pattern of lncRNA occupancy provides additional support for the idea that the lncRNAs explored here might have a role in recruiting protein complexes to promoter regions of target genes to regulate transcription, either by activating the formation of the pre-initiation complex or recruiting histone modifiers. However, while additional experiments are needed to confirm the roles of these nuclear lncRNAs in the parasite, using ChIRP-seq, we demonstrate that genome-wide collections of RNA binding sites can be used to discover the DNA sequence motifs enriched by lncRNAs. These findings signify the existence of lncRNA target sites in the genome, an entirely new class of regulatory elements that could be essential for transcriptional regulation in the malaria parasite.

Genetic disruption of lncRNA-ch14, a transcript detected specifically in gametocytes, demonstrates that this lncRNA plays an important role in sexual differentiation and is required for onward transmission to the mosquito **(Fig. 7a and b)**. This finding is supported by our transcriptomic analysis where we identified significant downregulation of genes involved in sexual differentiation including NEK and MAP kinases^74,75^ ookinete/oocyst development^76^ and microtubule function (i.e. dyneins and kinesins) most likely important in reshaping the parasite into sexual stages **(Fig. 7f and 7g, Fig. S7)**. Importantly, the skewed sex ratio of the LlncRNA-ch14 parasites does not completely account for the dramatic decrease in the ability of these parasites to be transmitted to mosquitoes. Indeed, our data suggest that the gametocytes of the LlncRNA-ch14 parasites are less infectious, a result that is also supported by the decrease in exflagellation of the male gametocytes **(Fig. 7c)**. It is currently difficult to assess infectiousness of female gametocytes, but we would hypothesize that these are also impacted by the disruption of lncRNA-ch14. While further experiments will be needed to further validate the effect of the full deletion or downregulation of the lncRNA-ch14 transcript, the results presented here confirm that some of the lncRNAs identified in this study play a role in the parasite’s sexual development and onward transmission to the mosquito.

Compared to the progress made in understanding lncRNA biology in higher eukaryotes, the field of lncRNAs in *Plasmodium* is still evolving. Analysis of promoter and gene body regions with available histone modification datasets (H3K9me3, H3K36me3, and H3K9ac) are still needed for further annotation of these candidate lncRNAs. It is clear that lncRNAs represent a new paradigm in chromatin remodeling and genome regulation. Therefore, this newly generated dataset will not only assist future lncRNA studies in the malaria parasite but will also help in identifying parasite-specific gene expression regulators that can ultimately be used as new anti-malarial drug targets.

## Materials and Methods

### Parasite culture

*P. falciparum* 3D7 strain at ∼ 8% parasitemia was cultured in human erythrocytes at 5% hematocrit in 25 mL of culture as previously described in^77^. Two synchronization steps were performed with 5% D-sorbitol treatments at ring stage within eight hours. Parasites were collected at early ring, early trophozoite, and late schizont stages. Parasite developmental stages were assessed using Giemsa-stained blood smears.

### Nuclear and cytosolic RNA isolation

Highly synchronized parasites were first extracted using 0.15% saponin solution followed by centrifugation at 1500 x g for 10 mins at 4°C. Parasite pellets were then washed twice with ice cold PBS and re-collected at 1500 x g. Parasite pellets were resuspended in 500 uL ice cold Cell Fractionation Buffer (PARIS kit, ThermoFisher; AM1921) with 10 uL of RNAse Inhibitor (SUPERaseIn 20U/uL, Invitrogen; AM2694) and incubated on ice for 10 minutes. Samples were centrifuged at 500 x g for 5 mins at 4°C. After centrifugation, the supernatant containing the cytoplasmic fraction was collected. Nuclei were resuspended in 500 uL Cell fractionation buffer and 15 uL RNAse Inhibitor as described above. To obtain a more purified nuclear fraction, the pellet was syringed with a 26G inch needle five times. The sample was incubated on ice for 10 mins and centrifuged at 500 x g for 5 mins at 4°C. The nuclear pellet was resuspended in 500 uL of ice-cold Cell Disruption Buffer (PARIS kit, ThermoFisher; AM1921). For both cytoplasmic and nuclear fractions, RNA was isolated by adding 5 volumes of Trizol LS Reagent (Life Technologies, Carlsbad, CA, USA) followed by a 5 min incubation at 37°C. RNA was then isolated according to manufacturer’s instructions. DNA-free DNA removal kit (ThermoFisher; AM1906) was used to remove potential genomic DNA contamination according to manufacturer’s instruction, and the absence of genomic DNA was confirmed by performing a 40-cycle PCR on the PfAlba3 gene using 200 to 500 ng input RNA.

### mRNA isolation and library preparation

Messenger RNA was purified from total cytoplasmic and nuclear RNA samples using NEBNext Poly(A) nRNA Magnetic Isolation module (NEB; E7490S) with manufacturer’s instructions. Once mRNA was isolated, strand-specific RNA-seq libraries were prepared using NEBNext Ultra Directional RNA Library Prep Kit for Illumina (NEB; E7420S) with library amplification specifically modified to accommodate the high AT content of *P. falciparum* genome: libraries were amplified for a total of 12 PCR cycles (45 s at 98°C followed by 15 cycles of 15 s at 98°C, 30 s at 55°C, 30 s at 62°C], 5 min 62°C). Libraries were then sequenced on Illumina NExtSeq500 generating 75 bp paired-end sequence reads.

### Sequence mapping

After sequencing, the quality of raw reads was analyzed using FastQC (https://www.bioinformatics.babraham.ac.uk/projects/fastqc/). The first 15 bases and the last base were trimmed. Contaminating adaptor reads, reads that were unpaired, bases below 28 that contained Ns, and reads shorter than 18 bases were also filtered using Sickle (https://github.com/najoshi/sickle)^78^. All trimmed reads were then mapped to the P. falciparum genome (v34) using HISAT2^79^ with the following parameters: –t, -- downstream-transcriptome assembly, --max-intronlen 3000, --no-discordant, -- summary-file, --known-splicesite-infile, --rna-strandness RF, and --novel-splicesite-outfile. After mapping, we removed all reads that were not uniquely mapped, not properly paired (samtools v 0.1.19-44428cd^80^) and are likely to be PCR duplicates (Picard tools v1.78, broadinstitute.github.io/picard/). The final number of working reads for each library is listed in **Supplemental Table S1**. For genome browser tracks, read coverage per nucleotide was first determined using BEDTools^81^ and normalized per million mapped reads.

### Transcriptome assembly and lncRNA identification

To identify lncRNAs in the nuclear and cytoplasmic fractions, we first merged all nuclear libraries and cytoplasmic libraries for each replicate, resulting in one pair of nuclear and cytoplasmic dataset per replicates. Next, we assembled the transcriptome (cufflinks v2.1.1)^82^ for each of the datasets using the following parameters: -p 8 -b PlasmoDB-34_Pfalciparum3D7_Genome.fasta -M PlasmoDB-34_Pfalciparum3D7.gff --library-type first strand -I 5000. After transcriptome assembly, we filtered out transcripts that are less than 200 basepairs and were predicted to be protein-coding (CPAT, http://lilab.research.bcm.edu). We then merge transcripts in both replicates using cuffmerge and removed any transcripts that are on the same strand and have more than 30% overlap with annotated regions (BEDTools intersect). Lastly, we further selected transcripts that have both primary and steady-state transcriptional evidence. For primary transcription, we used GRO-seq dataset (GSE85478) and removed any transcript that has a read coverage below 15% of the median expression of protein-encoding genes, as well as transcript that has an FPKM count less than 10 at any given stage. The same filtering criteria were also applied to the steady-state RNA-seq expression profiles.

To estimate the cellular location for each predicted lncRNA, we first calculated the summed read count of all nuclear libraries and the summed read count of all cytoplasmic libraries. Then, we measured the log2-fold change (log2FC) of the summed nuclear signal to the summed cytoplasmic signal. Any transcripts with a log2FC value above 0.5 were classified as nuclear enriched lncRNAs, and any transcripts with a log2 FC value below -0.5 were classified as cytoplasmic enriched lncRNAs. In addition, lncRNAs with a log2 FC value between the above thresholds were classified as lncRNAs expressed equally in both fractions.

### Western blot

Mixed-stage parasites were collected as described above. Parasite pellets were gently resuspended in 500 uL of ice-cold Cell Fractionation Buffer (PARIS kit, ThermoFisher; AM1921) and 50 uL of 10X EDTA-free Protease inhibitor (cOmplete Tablets, Mini EDTA-free, EASY pack, Roche; 05 892 791 001). Solution was incubated on ice for 10 mins and the sample was centrifuged for 5 mins at 4°C and 500 x g. The supernatant containing cytoplasmic fraction was collected carefully and the nuclear pellet was resuspended in 500 uL Cell Fractionation Buffer followed by needle-lysis 5x using 26 G inch needle. Nuclei were collected again at 4°C and 500 x g. The supernatant was discarded and the nuclei pellet in 500 uL of Cell Disruption Buffer (PARIS kit, ThermoFisher; AM1921) and incubated on ice for 10 minutes. The nuclear fraction was then sonicated 7x with 10 seconds on/30 seconds off using a probe sonicator. Extracted nuclear protein lysates were incubated for 10 mins at room temperature and centrifuged for 2 mins at 10,000 x g to remove cell debris. Seven micrograms of parasite cytoplasmic and nuclear protein lysates were diluted in a 2X laemmli buffer at a 1:1 ratio followed by heating at 95°C for 10 mins. Protein lysates are then loaded on an Any-KD SDS-PAGE gel (Bio-rad) and run for 1 hour at 125 V. Proteins were transferred to a PVDF membrane for 1 hr at 18 V, then stained using commercial antibodies generated against histone H3 (1:3,000 dilution, Abcam; ab1791) and PfAldolase (1:1,000 dilution, Abcam; ab207494), and secondary antibody, Goat Anti-Rabbit IgG HRP Conjugate (1:25,000 dilution, Bio-Rad; 1706515). Membranes were visualized using the Bio-Rad ChemidDoc MP Gel Imager.

### PiggyBac insertion analysis

To analyze lncRNA essentiality, we used piggyBac insertion sites from^39^, who performed saturation mutagenesis to uncover essential genes in *P. falciparum*. Since that study used an NF54 reference genome, we converted the coordinates to be applicable to the 3D7 reference genome (v38, PlasmoDB), using liftOver (Kent tools, UCSC Genome Bioinformatics group, https://github.com/ucscGenomeBrowser/kent). A chain file for the two genomes, needed for liftOver, was manually constructed as described here: http://genomewiki.ucsc.edu/index.php/LiftOver_Howto.

Custom Python scripts were used to overlap insertion site coordinates with lncRNA ranges to count the number of insertions that occurred in each lncRNA, as well as to locate TTAA sites (sites where piggyBac insertions could potentially occur) in the genome and count the number of TTAA sites in each lncRNA. These scripts were also used to determine the normalized location of each TTAA site and insertion site, in one of 50 windows either across the lncRNA range or also including the 5’ and 3’ flanking regions, which were each given 50% of the length of the lncRNA. The ratio of number of insertion sites to number of TTAA sites within a lncRNA was used as a loose measure of essentiality.

### Estimation of transcript stability

Read coverage values were calculated from total steady-state mRNA datasets (SRP026367, SRS417027, SRS417268, SRS417269) using BEDTools v2.25.0. The read counts were then normalized as described in the original publication, and ratios between RNA-seq and GRO-seq coverage values were calculated for each lncRNA and gene. This ratio reflects the relative abundance of the mature RNA transcript over its corresponding primary transcript and is a simple but convenient measurement for transcript stability.

### Reverse transcription PCR

Total RNA was isolated from 10 mL of mixed-stage asexual *P. falciparum* culture and 25 mL of late gametocyte stage culture. Total RNA quality was checked on an agarose gel and genomic DNA contamination was removed using a DNA-free DNA removal kit (ThermoFisher; AM1906) according to manufacturer’s instructions. The absence of genomic DNA was validated using a primer set targeting an intergenic region within PfAlba3 (PF3D7_1006200). Approximately 1 μg of DNase I treated RNA from each sample was used in a 35-cycle PCR reaction to confirm the absence of genomic DNA contamination. DNase-treated total RNA was then mixed with 0.1 μg of random hexamers, 0.6 μg of oligo-dT (20), and 2 μL 10 mM dNTP mix (Life Technologies) in total volume of 10 μL, incubated for 10 minutes at 70°C and then chilled on ice for 5 minutes. This mixture was added to a solution containing 4 μL 10X RT buffer, 8 μL 20 mM MgCl2, 4 μL 0.1 M DTT, 2 μL 20U/μl SuperaseIn and 1 μL 200 U/μL SuperScript III Reverse Transcriptase (Life Technologies). First-strand cDNA was synthesized by incubating the sample for 10 minutes at 25°C, 50 minutes at 50°C, and finally 5 minutes at 85°C. First strand cDNA is then mixed with 70 μL of nuclease free water, 30 μL 5x second-strand buffer (Life Technologies), 3 μL 10 mM dNTP mix (Life Technologies), 4 μL 10 U/μl *E. coli* DNA Polymerase (NEB), 1 μL 10 U/μL *E. coli* DNA ligase (NEB) and 1 μL 2 U/μL *E. coli* RNase H (Life Technologies). Samples were incubated for 2 h at 16°C and double stranded cDNA was purified using AMPure XP beads (Beckman Coulter). For testing transcription activity of predicted genes, 450 ng of double stranded cDNA was mixed with 10 pmole of both forward and reverse primers. DNA was incubated for 5 minutes at 95°C, then 30s at 98°C, 30s at 55°C, 30s at 62°C for 25 cycles. All primers used for PCR validation are listed in Supplemental Table S1.

### Single-cell sequencing and data processing

*P. falciparum* strain NF54 was cultured in O+ blood in complete RPMI 1640 culture medium at 37°C in a gas mixture of 5% O2/5% CO2/90% N2, as described previously 3, 78] Sexual commitment was induced at 1% parasitemia and 3% hematocrit and culture media were supplemented with 10% human serum. After 4, 6 and 10 days post sexual commitment, samples were taken from the culture for single cell sequencing. Cells from each day were loaded into separate inlets in a 10X chromium controller using the manufacturer’s instructions for a 10,000-target cell capture. Libraries for the days 4 and 6 samples were obtained using Chromium 10X version 2 chemistry, whereas libraries for the day 10 sample were obtained using version 3 chemistry. Cells were sequenced on a single lane of a HighSeq4000 using 150-bp paired-end reads. Raw reads were mapped to a custom gtf containing lncRNA coordinates appended to the *P. falciparum* 3D7 V3 reference genome (www.sanger.ac.uk/resources/downloads/protozoa/). Read mapping, deconvolution of cell barcodes and UMIs and the generation of single cell expression matrices were performed using the CellRanger pipeline v 3.0.0. LncRNA regions were labeled as ‘protein coding’ to be prioritized in STAR mapping in CellRanger. CellRanger was also run separately for each sample using the 3D7 reference genome that did not contain the appended non-coding regions for comparison. Resultant count matrices were loaded into the R package Seurat (v3.2.2) for pre-processing.

### Quality control and lncRNA expression

Single-cell transcriptomes (SCTs) were log-normalized, and expression scaled using Seurat (v.3.2.2). Each cell was assigned a stage by mapping to the Malaria Cell Atlas^83^ using scmap-cell (v1.8.0). Cells were assigned the stage of their closest neighbor in the Malaria Cell Atlas if they reached a cosine similarity of > 0.2. Cells identified as an early/late ring or late schizont containing < 50 UMI/cell and < 50 genes/cell were removed due to poor quality. Cells mapped to late stages, or cells not assigned to a stage in the Malaria Cell Atlas were removed if they contained < 100 UMIs/cell or < 80 genes/cell. Data from days 4, 6 and 10 were integrated together using Seurat’s IntegrateData function using 2000 integration anchors and 10 significant principal components. A variance stabilizing transformation was performed on the integrated matrix to identify the 750 most highly variable coding genes, and these were used to perform a principal component (PC) analysis. Significant PCs were then used to calculate three-dimensional UMAP embeddings using only coding genes. LncRNA expression was visualized on the UMAP embedding generated from coding gene expression using the package ggplot2 to assign stage-specific expression for the lncRNA.

### RNA in situ hybridization (RNA-FISH)

RNA FISH was performed with slight modifications as described by Mancio-Silva, 2012 33] on mixed-stage asexual and gametocyte stage parasites. Antisense RNA probes for seven nuclear lncRNAs; -TARE4, -178, -13, -1494, -271, -4076 -ch9, -ch14 and two cytoplasmic lncRNAs; -267, -643, were labeled by in vitro transcription in the presence of fluorescein. RNA FISH was also performed using sense RNA probes as controls. Briefly, fixed and permeabilized parasites were incubated with RNA probes overnight at 37°C. Parasites were washed with 2x SSC three times for 15 mins each at 45°C followed by one wash with 1x PBS for 5 mins at room temperature. The slides were mounted in a Vectashield mounting medium with DAPI and visualized using the Olympus BX40 epifluorescence microscope.

### Chromatin isolation by RNA purification (ChIRP)

ChIRP-seq experiments were performed in duplicate for all nuclear lncRNAs, at the time point of highest lncRNA expression and lowest expression. Synchronized parasite cultures were collected and incubated in 0.15% saponin for 10 min on ice to lyse red blood cells. Parasites were centrifuged at 3234 x g for 10 mins at 4°C and subsequently washed three times with PBS by resuspending in cold PBS and centrifuging for 10 mins at 3234 x g at 4°C. Parasites were cross-linked for 15 mins at RT with 1% glutaraldehyde. Cross-linking was quenched by adding glycine to a final concentration of 0.125 M and incubating for 5 mins at 37°C. Parasites were centrifuged at 2500 x g for 5 mins at 4°C, washed three times with cold PBS and stored at -80°C.

To extract nuclei, parasite was first incubated on in nuclear extraction buffer (10 mM HEPES, 10 mM KCl, 0.1 mM EDTA, 0.1 mM EGTA, 1 mM DTT, 0.5 mM 4-(2-aminoethyl) benzenesulfonyl fluoride hydrochloride (AEBSF), EDTA-free protease inhibitor cocktail (Roche) and phosphatase inhibitor cocktail (Roche)) on ice. After 30 mins, Igepal CA-360 (Sigma-Aldrich) was added to a final concentration of 0.25% and needle sheared seven times by passing through a 26 G ½ needle. Parasite nuclei were centrifuged at 2500 x g for 20 mins at 4°C and resuspended in shearing buffer (0.1% SDS, 1 mM EDTA, 10 mM Tris–HCl pH 7.5, EDTA-free protease inhibitor cocktail and phosphatase inhibitor cocktail). Chromatin was fragmented using the Covaris UltraSonicator (S220) to obtain 100-500 bp DNA fragments with the following settings: 5% duty cycle, 140 intensity incident power, 200 cycles per burst. Sonicated samples were centrifuged for 10 mins at 17000 x g at 4°C to remove insoluble material.

Fragmented chromatin was precleared using Dynabeads MyOne Streptavidin T1 (Thermo Fisher) by incubating for 30 mins at 37°C to reduce non-specific background. Per ChIRP sample using 1 mL of lysate, 10 uL each was removed for the RNA input and DNA input, respectively. Each sample was diluted in 2x volume of hybridization buffer (750mM NaCl, 1% SDS, 50mM Tris-Cl pH 7.5, 1mM EDTA, 15% formamide, 0.0005x volume of AEBSF, 0.01x volume of Superase-in (Ambion) and 0.01x volume of protease inhibitor cocktail). ChIRP probes used for each lncRNA (see Supplemental Table S1) were pooled, heated at 85°C for 3 mins and cooled on ice. ChIRP probes were added to each sample (2 uL of 100 uM pooled probes per sample) and incubated at 37°C with end-to-end rotation for 4 hours. Prior to completion of hybridization, Dynabeads MyOne Streptavidin T1 beads were washed three times on a magnet stand using lysis buffer (50mM Tris-Cl pH 7, 10mM EDTA, 1% SDS). After the hybridization, 100 uL of washed T1 beads were added to each tube and incubated for 30 mins at 37°C. Beads were washed with wash buffer (2x SSC, 0.5% SDS, 0.005x volume of AEBSF) and split evenly for isolation of DNA and RNA fractions.

For RNA isolation, the RNA input and chromatin-bound beads were resuspended in RNA elution buffer (100mM NaCl, 10mM Tris-HCl pH 7.0, 1mM EDTA, 0.5% SDS, 1mg/mL Proteinase K), incubated at 50°C for 45 mins, boiled at 95°C for 15 mins and subjected to trizol:chloroform extraction. Genomic DNA contamination was removed using a DNA-free DNA removal kit (ThermoFisher; AM1906) according to manufacturer’s instructions. The absence of genomic DNA was validated using a primer set targeting an intergenic region within PfAlba3 (PF3D7_1006200) in a 35-cycle PCR reaction. DNase-treated RNA was then mixed with 0.1 μg of random hexamers, 0.6 μg of oligo-dT (20), and 2 μL 10 mM dNTP mix (Life Technologies) in total volume of 10 μL, incubated for 10 minutes at 70°C and then chilled on ice for 5 minutes. This mixture was added to a solution containing 4 μL 10X RT buffer, 8 μL 20 mM MgCl2, 4 μL 0.1 M DTT, 2 μL 20U/μl SuperaseIn and 1 μL 200 U/μL SuperScript III Reverse Transcriptase (Life Technologies). First-strand cDNA was synthesized by incubating the sample for 10 minutes at 25°C, 50 minutes at 50°C, and finally 5 minutes at 85°C followed by a 20 min incubation with 1 μL 2 U/μL *E. coli* RNase H (Life Technologies) at 37°C. Prepared cDNA was then subjected to quantitative reverse-transcription PCR for the detection of enriched TARE-4 and serine tRNA ligase transcripts with the following program: 5 minutes at 95°C, 30 cycles of 30s at 98°C, 30s at 55°C, 30s at 62°C and a final extension 5 min at 62°C. All primers used for PCR validation are listed in Supplemental Table S1.

Libraries from the ChIRP samples were prepared using the KAPA Library Preparation Kit (KAPA Biosystems). Libraries were amplified for a total of 12 PCR cycles (12 cycles of 15 s at 98°C, 30 s at 55°C, 30 s at 62°C]) using the KAPA HiFi HotStart Ready Mix (KAPA Biosystems). Libraries were sequenced with a NextSeq500 DNA sequencer (Illumina). Raw read quality was first analyzed using FastQC (https://www.bioinformatics.babraham.ac.uk/projects/fastqc/). Reads were mapped to the *P. falciparum* genome (v38, PlasmoDB) using *Bowtie2* (v2.4.2). Duplicate, unmapped, and low quality (MAPQ < 20) reads were filtered out using *Samtools* (v1.9), and only uniquely mapped reads were retained. All libraries, including the input, were then normalized by dividing by the number of mapped reads in each of them. For each nucleotide, the signal from the input library was then subtracted from each of the ChIRP-seq libraries, and any negative value was replaced with a zero. Genome tracks were generated by the R package *ggplot2*.

### Peak Calling

Peaks were called using *PePr*. For a given lncRNA of interest, the tool was run in differential binding analysis mode using the filtered ChIRP-seq libraries for when the lncRNA was active versus non-active with the following parameters specified: *-*peaktype broad *-*threshold 1e-10. The top 25% of all reported peaks were selected because they exhibited the strongest signal (see Supplementary Figure S4) and used in downstream analyses. For differential gene expression analysis, the closest gene to each peak was selected, and its expression in the two stages (active versus inactive) was obtained from37,84,85.

### LncRNA-ch14 gene disruption

Gene knockout (KO) for the long non-coding RNA 14 (lncRNA-14) spanning position (Ch14:3,148,960 - 3,150,115) on chromosome 14 was performed using a two-plasmid design. The plasmid pDC2-Cas9-sgRNA-hdhfr^86^, gifted from Marcus Lee (Wellcome Sanger Institute) contains the SpCas9, a site to express the sgRNA, and a positive selectable marker human dihydrofolate reductase (*hdhfr*). The sgRNA was selected from the database generated by^87^ and cloned into pDC2-Cas9-sgRNA-hdhfr at the BbsI restriction site. The homology directed repair plasmid (modified pDC2-donor-*bsd* without eGFP) was designed to insert a selectable marker, blasticidin S-deaminase *(bsd)*, disrupting the lncRNA-14 region. The target specifying homology arm sequences were isolated through PCR amplification and gel purification. The right homology regions (RHR), and the left homology regions (LHR) of each gene were then ligated into the pDC2-cam-Cas9 linear vector via Gibson assembly. The final donor vectors were confirmed by restriction digests and Sanger sequencing.

Plasmids were isolated from 250 mL cultures of *Escherichia coli* (XL10-Gold Ultracompetent Cells, Agilent Cat. 200314) and 60 μg of each plasmid was used to transfect ring stage parasites. 24-hrs before transfection, mature parasite cultures (6-8% parasitemia) were magnetically separated using magnetic columns (MACS LD columns, Miltenyi Biotec) and diluted to 1% parasitemia containing 0.5 mL fresh erythrocytes^88^. The next day, ∼3% ring stage parasites were pelleted and washed in 4 mL of cytomix^89^. 200 μl of the infected erythrocytes were resuspended with the two plasmids in cytomix to a total volume of 400 μl in a 0.2 cm cuvette. Electroporation was performed with a single pulse at 0.310 kV and 950 μF using the Biorad GenePulser electroporator. Cells were immediately transferred to a flask containing 12 mL media and 400 μl erythrocytes. The media was exchanged five hours post electroporation with 12 mL of fresh media. The following day, fresh culture media was added and supplemented with 2.5 nM WR99210 and 2.5 μg/mL blasticidin (RPI Corp B12150-0.1). Media and drug selection were replenished every 48 hours. After 14 days, the culture was split into two flasks and 50 μl of erythrocytes were added every two weeks. Once parasites were detected by microscopy, WR99210 was removed (selection for Cas9). Integration of the bsd gene was confirmed by gDNA extraction and PCR.

### Isolation of LncRNA14-KO clone

To generate genetically homogenous parasite lines, the transfected parasites were serially diluted to approximately 0.5%, into 96 well plates. 200 μl final volume of cultured parasites were incubated with bsd drug selection for 1 month with weekly erythrocytic and media changes for the first 2 weeks of dilution followed by media changes every 2 days until parasite recovery is observed through Giemsa-stained smears.

### Verification of LncRNA14-KO line

Genomic DNA (gDNA) was extracted and purified using DNeasy Blood &amp; Tissue kit (Qiagen) following instructions from the manufacturer. The diagnostic and genotyping PCR analysis was used to genotype the KO lines using three primer pairs Set A: BSD-F/Lnc14KO-R, Set B: Lnc14_F/Lnc14RT2-R and Set C: Lnc14_F/Lnc14_R annealing to the genomic sequence of the targeted locus. The PCR amplification was done using 2xKAPPA master mix for thirty cycles with an annealing temperature of 50°C and an extension temperature of 62°C. The PCR amplicons were analyzed on a 1% agarose gel electrophoresis.

For whole genome sequencing, genomic DNAs were fragmented using a Covaris S220 ultrasonicator and libraries were generated using KAPA LTP Library Preparation Kit (Roche, KK8230). To verify that the insertion was present in the genome at the correct location in both transfected lines, reads were mapped using Bowtie2 (version 2.4.4) to the *P. falciparum* 3D7 reference genome (v48, PlasmoDB), edited to include the insertion sequence in the intended location. IGV (Broad Institute) was used to verify that reads aligned to the insertion sequence.

### LncRNA14-KO line genome-wide sequencing and variant analysis

Libraries were sequenced using a NovaSeq 6000 DNA sequencer (Illumina), producing paired-end 100-bp reads. To verify that the insertion was present in the genome at the correct location in both transfected lines, reads were mapped using Bowtie2 (version 2.4.4) to the *P. falciparum* 3D7 reference genome (v48, PlasmoDB), edited to include the insertion sequence in the intended location. IGV (Broad Institute) was used to verify that reads aligned to the insertion sequence. To call variants (SNPs/indels) in the transfected lines compared to a previously sequenced NF54 control line, genomic DNA reads were first trimmed of adapters and aligned to the *Homo sapiens* genome (assembly GRCh38) to remove human-mapped reads. Remaining reads were aligned to the *P. falciparum* 3D7 genome using bwa (version 0.7.17) and PCR duplicates were removed using PicardTools (Broad Institute). GATK HaplotypeCaller (https://gatk.broadinstitute.org/hc/en-us) was used to call variants between the sample and the 3D7 reference genome for both the transfected lines and the NF54 control. Only variants that were present in both transfected lines but not the NF54 control line were kept. We examined only coding-region variants and removed those that were synonymous variants or were in *var, rifin*, or *stevor* genes. Quality control of variants was done by hard filtering using GATK guidelines.

### Assessment of gametocyte development

Viability of gametocytes was assessed via microscopy in parasite laboratory strains NF54 and two of the KO clones, F2 and B1. The morphology of parasite gametocytes was assessed in a Giemsa-stained thin blood smear. Gametocytes were classified either as viable (normal intact morphology of mature gametocytes) or dead (deformed cells with a decrease in width, a thin needle-like appearance or degraded cytoplasmic content).

### Gametocyte cultures and mosquito feeding

This was performed as outlined previously 50]. Briefly, asexual stage cultures were grown in RPMI-1640 containing 2 mM L-glutamine, 50 mg/L hypoxanthine, 25 mM HEPES, 0.225% NaHCO_3_, contained 10% *v/v* human serum in 4% human erythrocytes. Five mL of an asexual stage culture at 5% parasitaemia was centrifuged at 500 × *g* for 5 min at room temperature. Gametocyte cultures were initiated at 0.5% asynchronous asexual parasitemia from low passage stock and maintained up to day 18 with daily media changes but without any addition of fresh erythrocytes. The culture medium was changed daily for 15-18 days, by carefully aspirating ∼ 70-80% of the supernatant medium to avoid removing cells, and 5 mL of fresh complete culture medium was added to each well. Giemsa-stained blood smears were made every alternate day to confirm that the parasites remained viable. Instead of a gas incubator, cultures were maintained at 37°C in a candle jar made of glass desiccators. On day 15 to 18, gametocyte culture, containing largely mature gametocytes, were used for mosquito feeds. Cultures were transferred to pre-warmed tubes and centrifuged at 500 x *g* for 5 min. The cells were diluted in a pre-warmed 50:50 mixture of uninfected erythrocytes and normal human serum to achieve a mature gametocytemia of 0.2% and the resulting ‘feeding mixture’ was placed into a pre-warmed glass feeder. Uninfected *Anopheles stephensi* mosquitoes, starved overnight of sugar water, were allowed to feed on the culture for 30 min. Unfed mosquitoes were removed, and the mosquito cups were placed in a humidified 26°C incubator, with 10% sugar-soaked cotton pads placed on top of the mosquito cage.

### Oocyst and salivary gland sporozoite quantification

On days 11 and 17 after the infective-blood meal, mosquitoes were dissected and midguts or salivary glands, respectively, were harvested for sporozoite counts. Day 11 midguts were stained with mercurochrome and photographed for oocyst counts by brightfield and phase microscopy using an upright Nikon E600 microscope with a PlanApo 4× objective. On day 17, salivary glands from ∼20 mosquitoes were pooled, homogenized, and released sporozoites were counted using a haemocytometer.

### Gametocyte quantification, sex determination and exflagellation assay

Between days 15 to 18, blood smears were prepared from gametocyte cultures, fixed with methanol and stained with Giemsa (Sigma GS500), prepared as a 1:5 dilution in buffer (pH=7.2) made using Gurr buffered tablets (VWR #331942F) and filtered. Slides were stained for 20 min, washed with buffer, and allowed to dry before observation using a Nikon E600 microscope with a PlanApo 100× oil objective. For calculation of gametocytemias and male: female ratios, at least 500 mature gametocytes were scored per slide. To count exflagellation centers, 500 μl of mature gametocyte culture was centrifuged at 500g for four min and the resulting pellet was resuspended in equal volume of prewarmed normal human serum. Temperature was dropped to room temperature to activate gametogenesis and after 15 min incubation 10μl of culture was transferred to a glass slide and covered with a cover slip. Exflagellation centers were counted at 40x objective in at-least ten fields for each gametocyte culture. To avoid bias, microscopic examination was performed in a blinded fashion by a trained reader.

### LncRNA14-KO transcriptome analysis

Libraries were prepared from the extracted total RNA, first by isolating mRNA using the NEBNext Poly(A) mRNA Magnetic Isolation Module (NEB), then using the NEBNext Ultra Directional RNA Library Prep Kit (NEB). Libraries were amplified for a total of 12 PCR cycles (12 cycles of 15 s at 98°C, 30 s at 55°C, 30 s at 62°C]) using the KAPA HiFi HotStart Ready Mix (KAPA Biosystems). Libraries were sequenced using a NovaSeq 6000 DNA sequencer (Illumina), producing paired-end 100-bp reads.

FastQC (https://www.bioinformatics.babraham.ac.uk/projects/fastqc/), was used to assess raw read quality and characteristics, and based on this information, the first 11 bp of each read and any adapter sequences were removed using Trimmomatic (http://www.usadellab.org/cms/?page=trimmomatic). Bases were trimmed from reads using Sickle with a Phred quality threshold of 20 (https://github.com/najoshi/sickle). These reads were mapped against the *Homo sapiens* genome (assembly GRCh38) using Bowtie2 (version 2.3.4.1) and mapped reads were removed. The remaining reads were mapped against the *Plasmodium falciparum* 3D7 genome (v48, PlasmoDB) using HISAT2 (version 2.2.1), using default parameters. Uniquely mapped, properly paired reads with mapping quality 40 or higher were retained using SAMtools (http://samtools.sourceforge.net/), and PCR duplicates were removed using PicardTools (Broad Institute). Genome browser tracks were generated and viewed using the Integrative Genomic Viewer (IGV) (Broad Institute).

Raw read counts were determined for each gene in the *P. falciparum* genome using BedTools (https://bedtools.readthedocs.io/en/latest/#). to intersect the aligned reads with the genome annotation. Differential expression analysis was done by use of R package DESeq2 (https://bioconductor.org/packages/release/bioc/html/DESeq2.html) to call up- and down-regulated genes with an adjusted P-value cutoff of 0.05. Gene ontology enrichment was done using PlasmoDB (https://plasmodb.org/plasmo/app). Volcano plots were generated using R package EnhancedVolcano (https://bioconductor.org/packages/release/bioc/html/EnhancedVolcano.html).

### Statistical analysis

Descriptive statistics were calculated with GraphPad Prism version 9.1.2 (GraphPad Software, San Diego, CA, USA) for determining mean, percentages, standard deviation and plotting of graphs. Excel 2013 and GraphPad Prism 9.1.2 were used for the calculation of gametocytemia of microscopic data.

## Figure Legends

**Supplementary Fig. S1:** Spearman correlation among gene expression levels of nuclear fraction, cytoplasmic fraction, and steady-state mRNA across cell cycle of *P. falciparum*.

**Supplementary Fig. S2**: Essentiality of lncRNAs in *P. falciparum*. (a) Number of piggyBac insertions per possible insertion site (TTAA site) for telomeric, subtelomeric, and other lncRNAs. Telomeric lncRNAs have significantly fewest insertions/TTAA sites, followed by subtelomeric lncRNAs. p-value were calculated using the *t*-*test*. (b) Plot showing normalized density of piggyBac insertions and TTAA sites across all detected lncRNAs including flanking regions for each lncRNA, representing 50% of the lncRNA length on both the 5’ and 3’ sides.

**Supplementary Fig. S3**. (a) Total RNA was extracted from both asexual and gametocyte stage parasites. RNA quality was validated on agarose gel. (b) Genomic DNA was removed and verified using reverse transcription polymerase chain reaction (RT-PCR) with primers designed to amplify a fragment of *pfAlba3* gene (PF3D7_1006200). Primers were designed on both sides of intron 1, yielding a 429 bp PCR product from genomic DNA and a 164 bp PCR product from cDNA. The absence of PCR product amplified from RNA confirms the absence of gDNA contamination. (c) RT-PCR validation of two selected lncRNA that are most abundantly expressed at the gametocyte stage with high level of H3K9me3 mark.

**Supplementary Fig. S4**. Genome wide ChIRP-seq signal for lncRNAs. All lncRNA libraries, including the input, were normalized by dividing by the numbers of mapped reads in each of them. For each nucleotide, the signal from the input library was then subtracted from each of the ChIRP-seq libraries, and any negative value was replaced with a zero. Genome tracks are displayed for each chromosome.

**Supplementary Fig. S5**. Average of ChIRP-seq signals across all 5’ and 3’ UTRs and gene bodies. For a given lncRNA of interest we examined the regions of the genome that its corresponding peaks overlap. We extended a 1000bp window from the 5’ and 3’ ends of the nearest gene to each peak. We aggregated these genomic windows together with the gene body (normalized to 1000bp to account for different gene lengths) into a continuous 3000b window (x-axis). For each base position we plotted the density (the number of peaks overlapping the respective position divided by the total number of peaks) on the y-axis.

**Supplementary Fig. S6**. (a) LncRNA-ch14 disruption strategy using the CRISPR-Cas9 tool through BSD^r^ insertion. (b) PCR Gel verification for lncRNA-ch14 disruption through PCR of 5’Arm insertion (Left), BSD^r^ and 3’Arm insertion (Middle) and entire lncRNA-ch14 segment (Right).

**Supplemental Fig. S7**. Gene expression profiles between WT and △lncRNA-ch14 lines are presented in bar graph by Log10P (y-axis) in asexual stage mature parasites (Left) and mature gametocytes (Right).

**Supplementary Table S1**. Putative lncRNAs identified and investigated in this study. Chromosome, coordinates, and expression profile for all lncRNAs identified along with probe and primer sequences used to investigate several stage-specific lncRNAs.

**Table S1a:** Mapping statistic for Nuclear and Cytoplasmic RNA-seq data.

**Table S1b:** Putative lncRNAs identified in this study.

**Table S1c:** LncRNAs used for ChIRP-seq analysis.

**Table S1d:** ChIRP-seq probes and sequences used for identifying lncRNA-chromatin interactions.

**Table S1e:** Sequences of primers used for RT-PCR and the generation of RNA-FISH probes.

**Table S1f:** Peak calls for ChIRP-seq against lncRNA-TARE

**Table S1g:** Peak calls for ChIRP-seq against lncRNA-13

**Table S1h:** Peak calls for ChIRP-seq against lncRNA-178

**Table S1i:** Peak calls for ChIRP-seq against lncRNA-271

**Table S1j:** Peak calls for ChIRP-seq against lncRNA-1494

**Table S1k:** Peak calls for ChIRP-seq against lncRNA-ch9

**Table S1l:** Peak calls for ChIRP-seq against lncRNA-ch14

**Supplementary Table S2**. Whole genome sequencing results showing identified mutations in NF54 control and △lncRNA-ch14 transfected lines. The mutation in the *gdv1* gene essential for gametocyte formation exists only in the parental NF54 parasites highlighted in red.

## Acknowledgement

This work was supported by NIH grants to KGLR (nos. 1R01 AI136511 and R21 AI142506-01) and by the University of California, Riverside to KGLR (no. NIFA-Hatch-225935). We also thank the parasitology and insectary core facilities at the JHMRI and Bloomberg Philanthropies for their support of these facilities.

## Author contributions

Conceptualization was the responsibility of KGLR that also supervised the project together with ML, WSN and PS. Methodology was performed by GB, XML, SA, ZC, DW, TH, TW, AC, JP, AKT, GU, JC, TT and SD; GB, XML, SA, BH and KGLR carried out the analysis. Software, formal analysis, and data curation were provided by SA, BH and SD. GB, ZC, SA and KGLR wrote the original draft. All authors reviewed and edited the manuscript.

