## Supplementary material for "Deciphering the non-coding code of pathogenicity and sexual differentiation in the human malaria parasite": Supp Figs

Supplementary Fig. S1

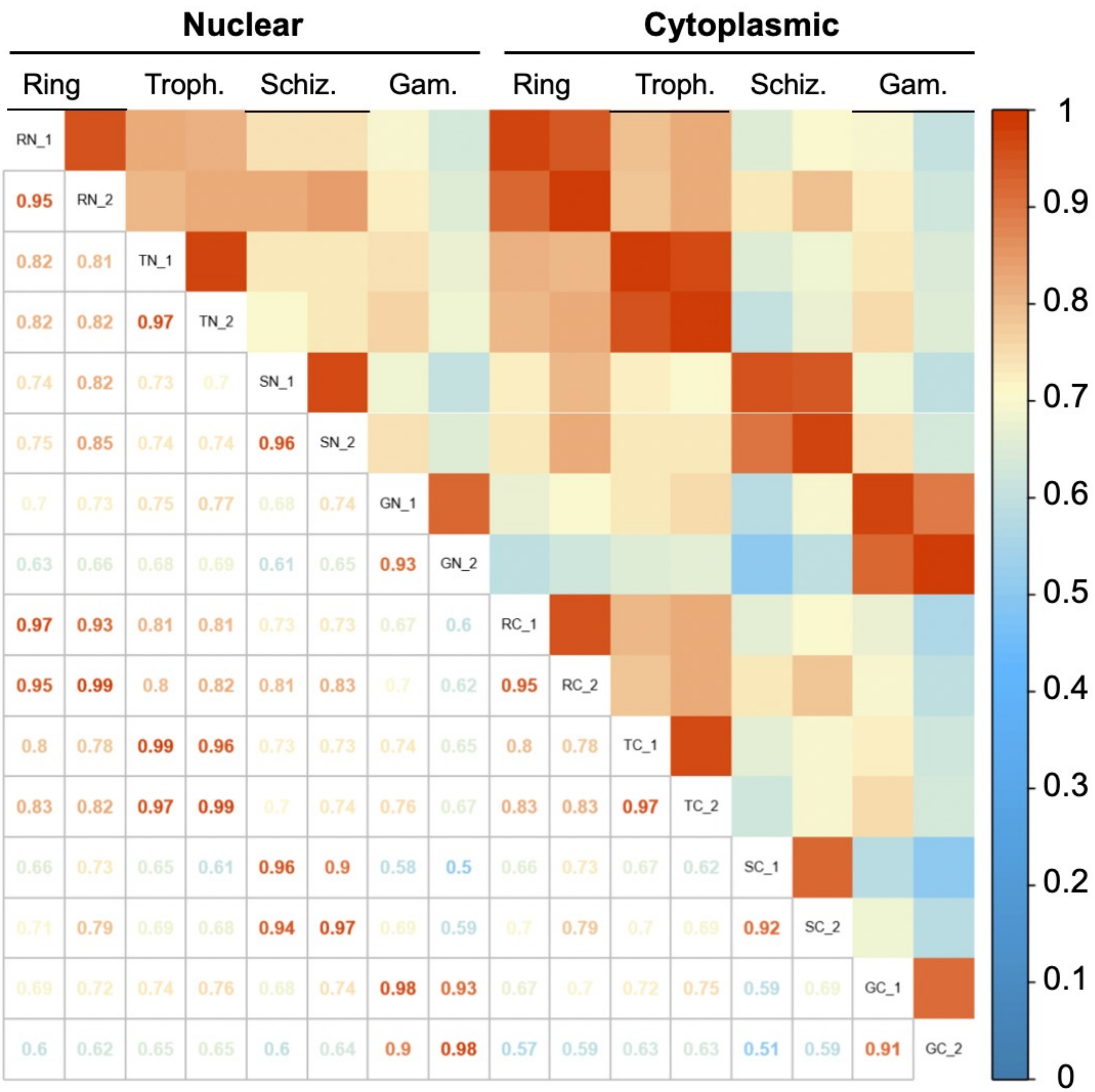

Supplementary Fig. S2

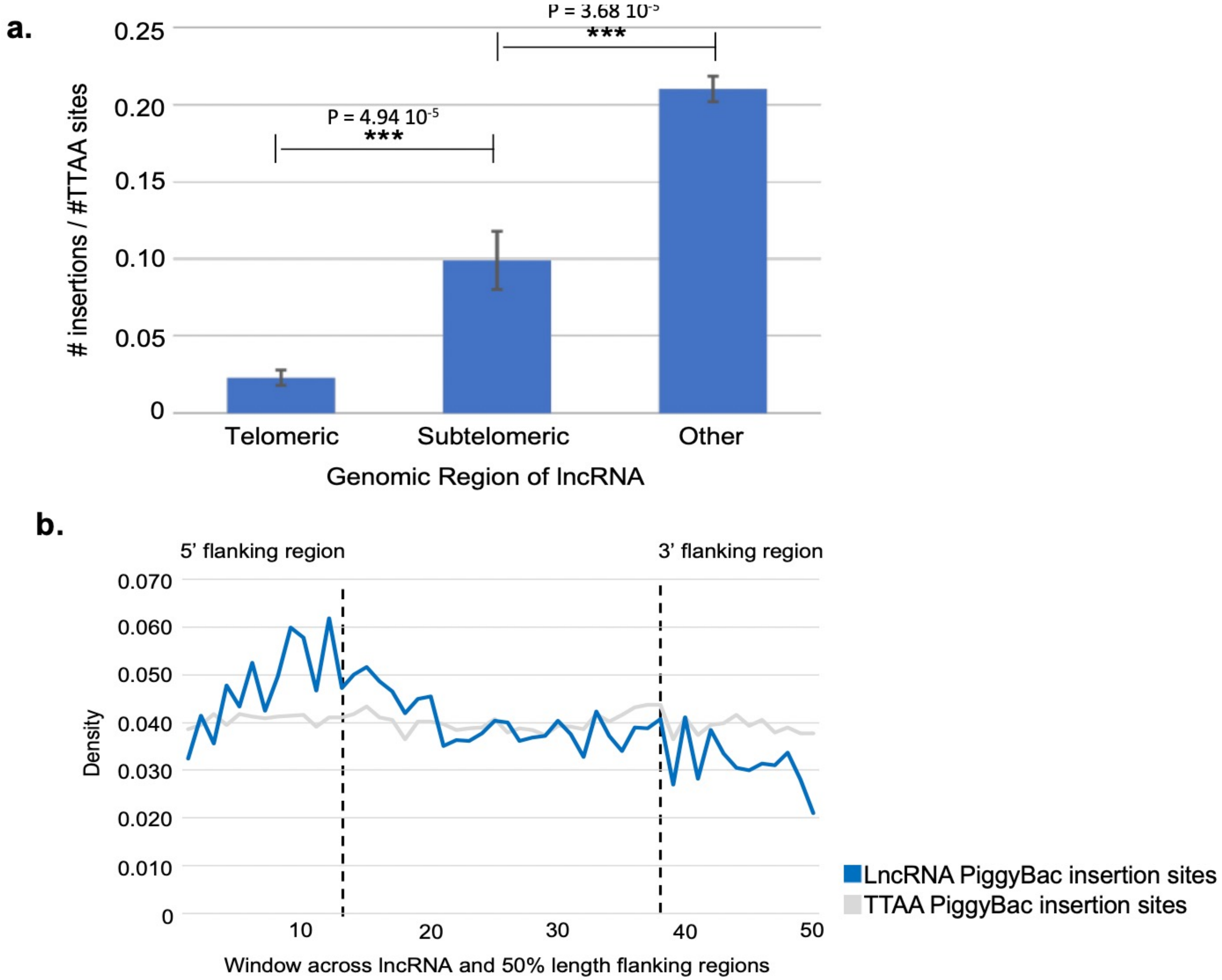

Supplementary Fig. S3

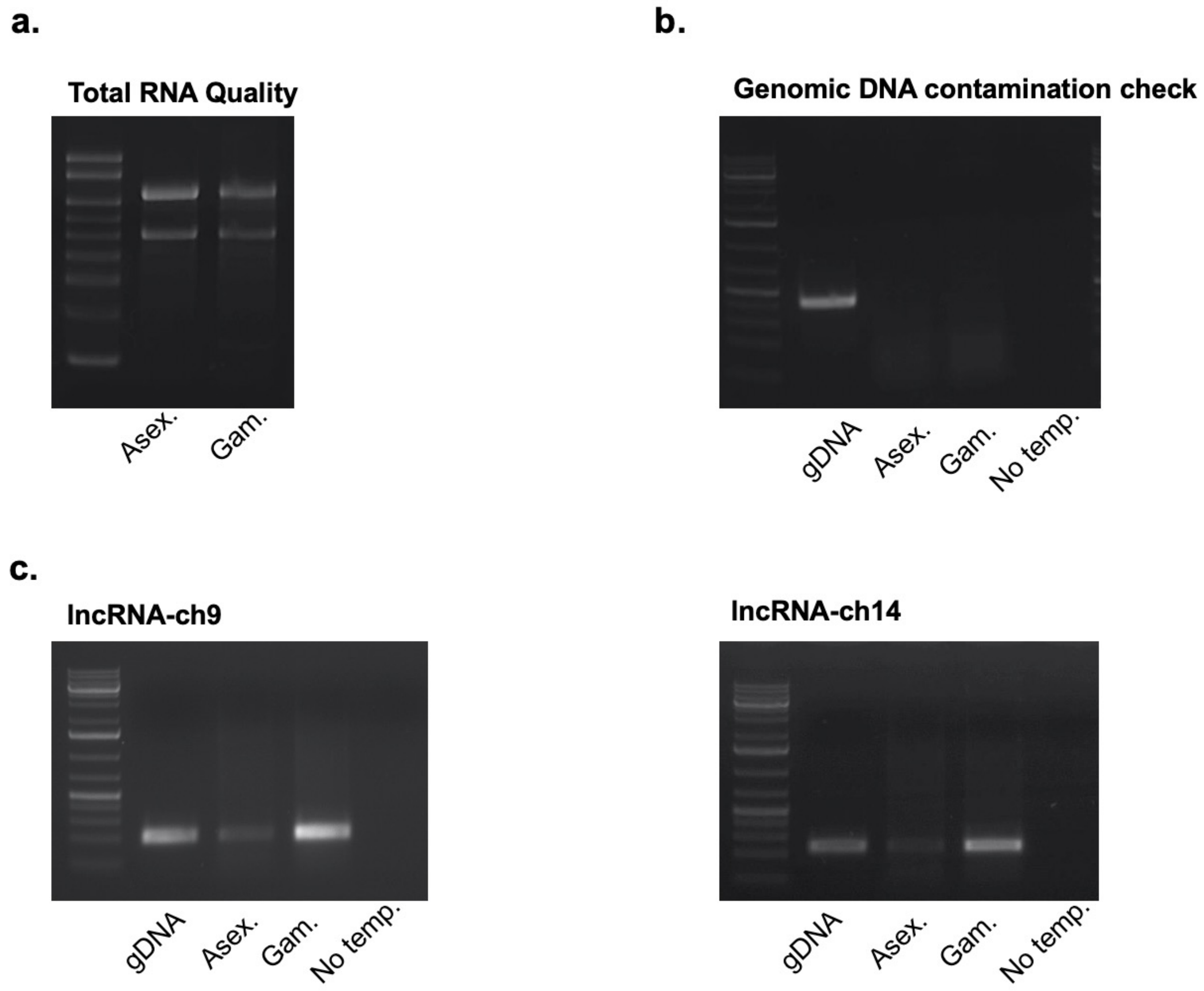

Supplementary Fig. S4

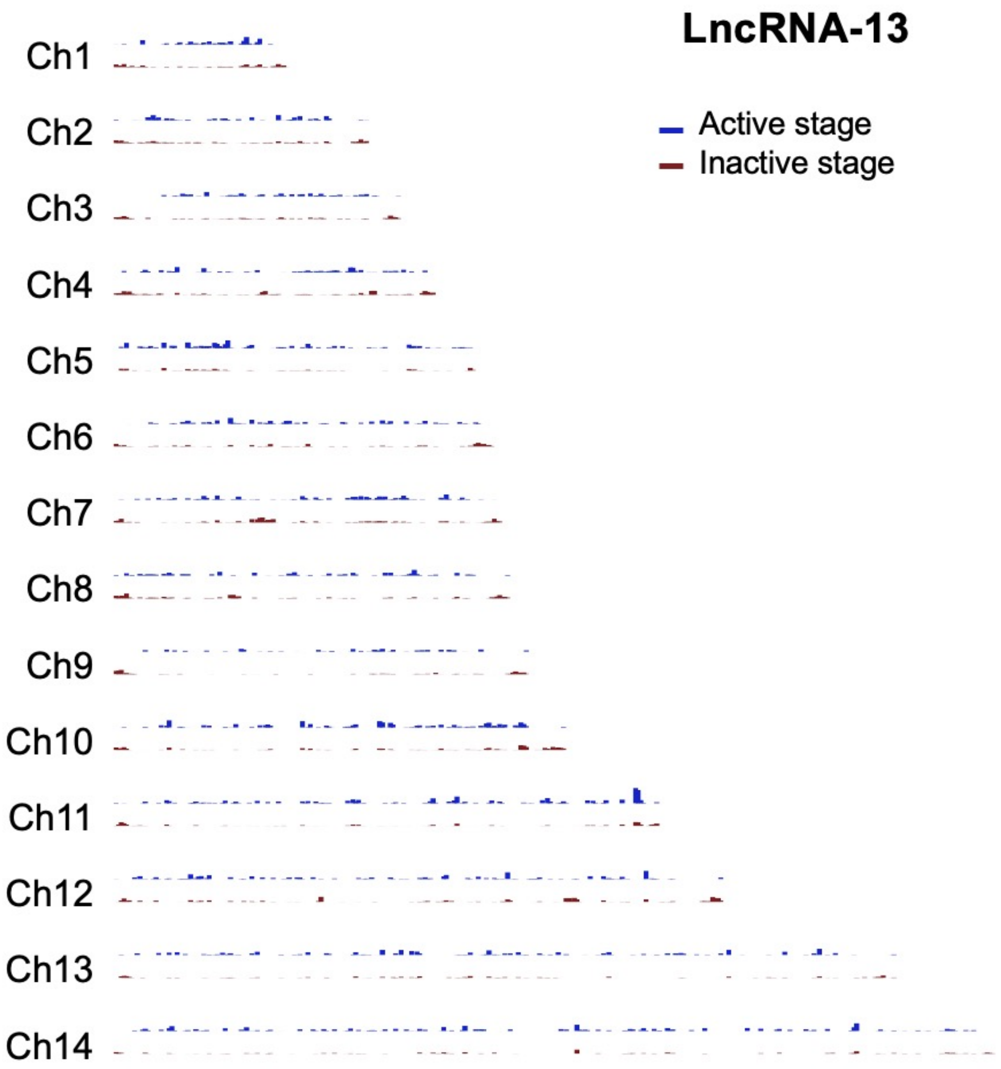

Supplementary Fig. S4

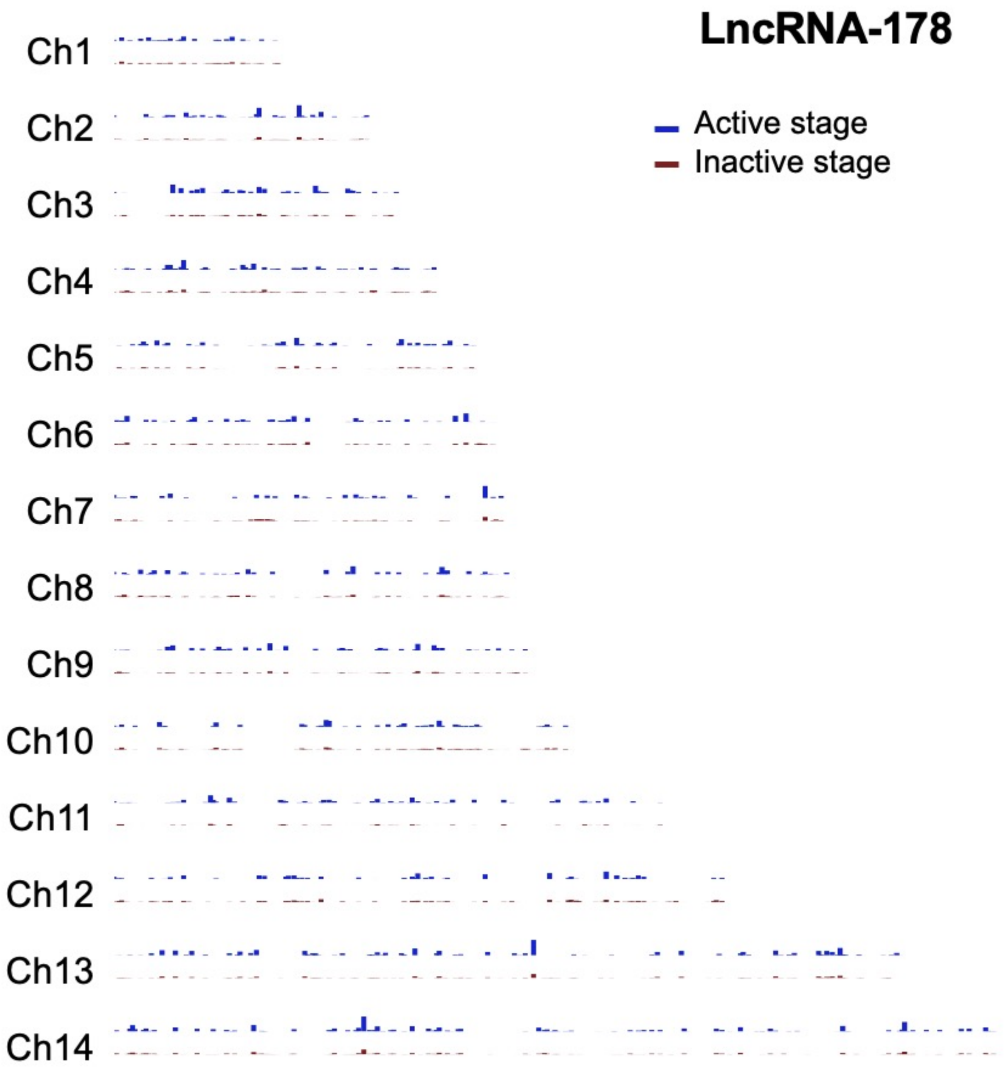

Supplementary Fig. S4

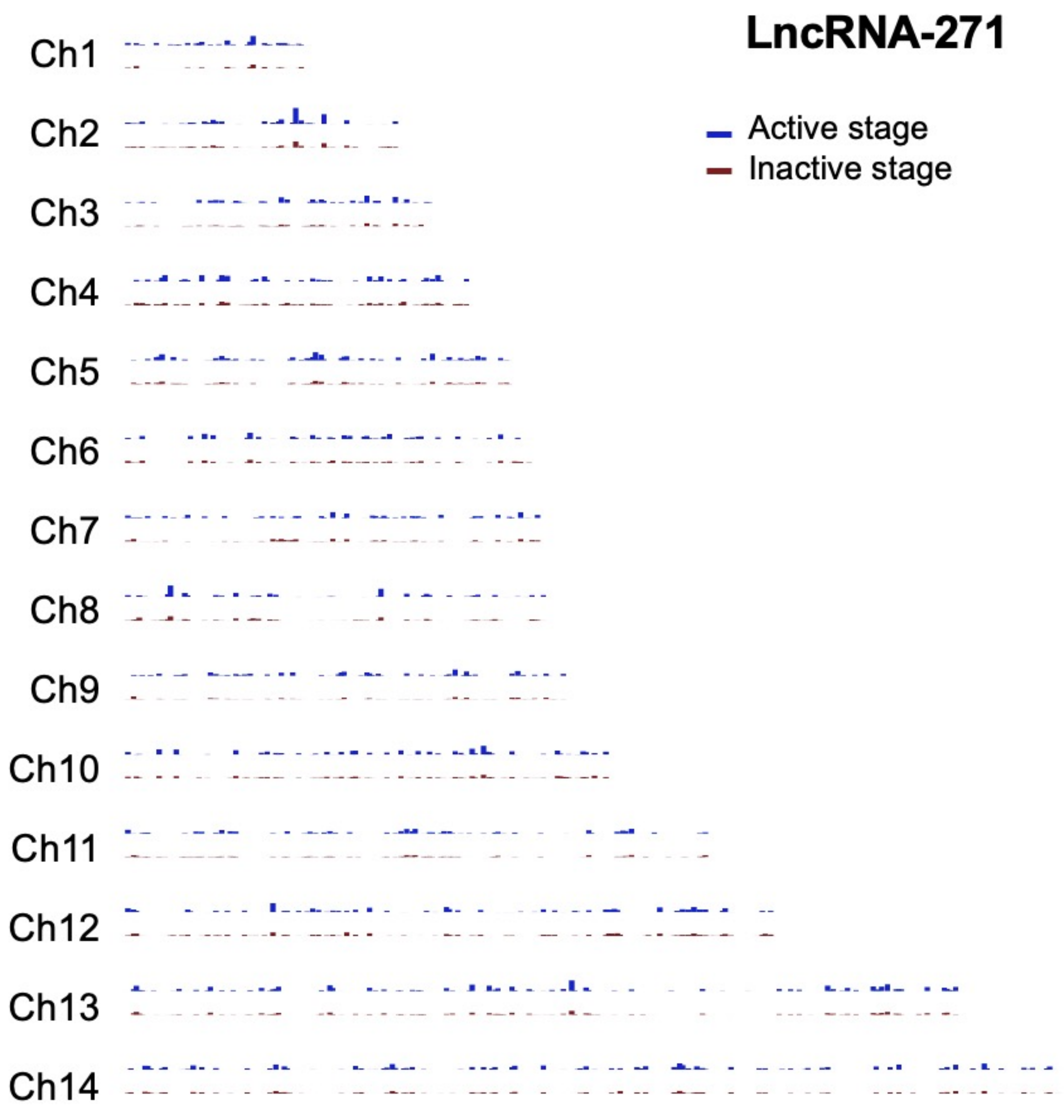

Supplementary Fig. S4

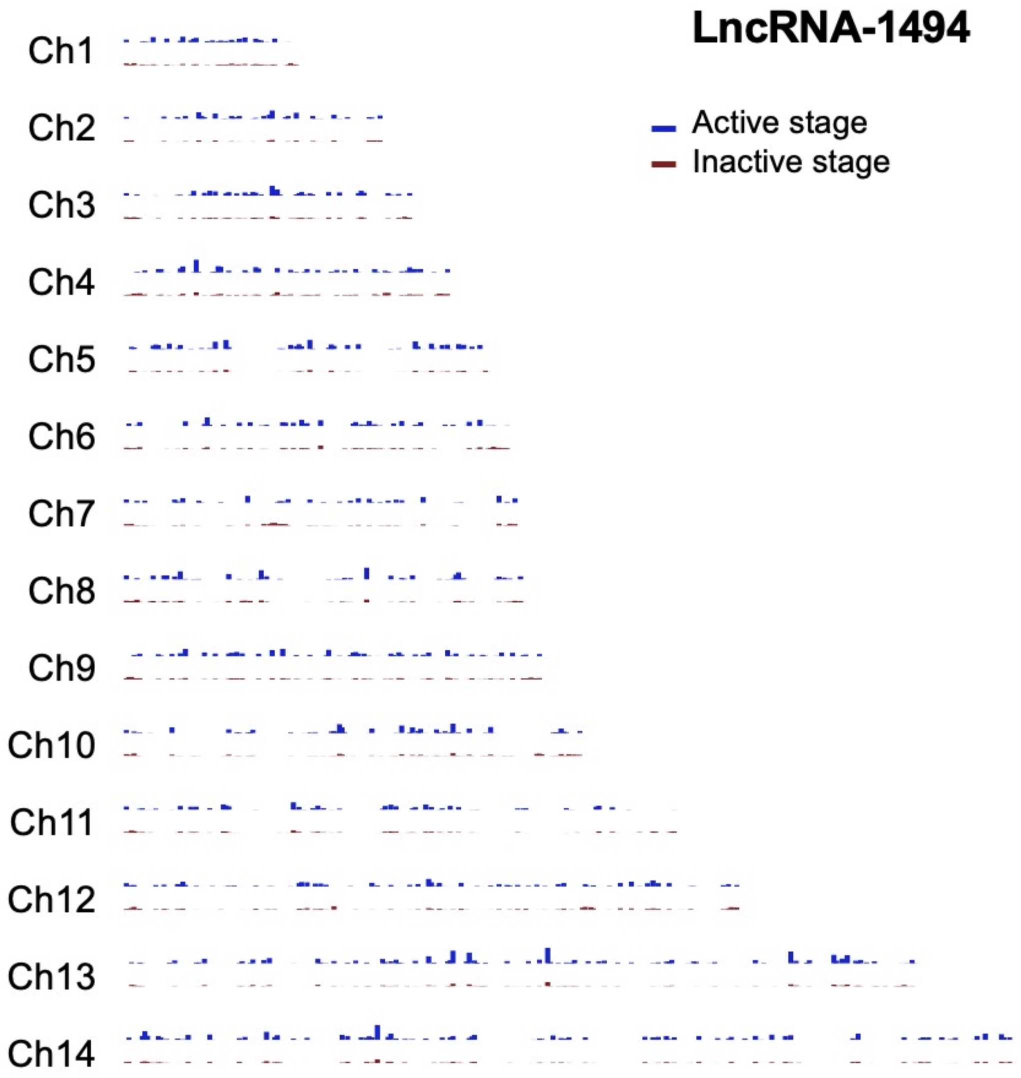

Supplementary Fig. S4

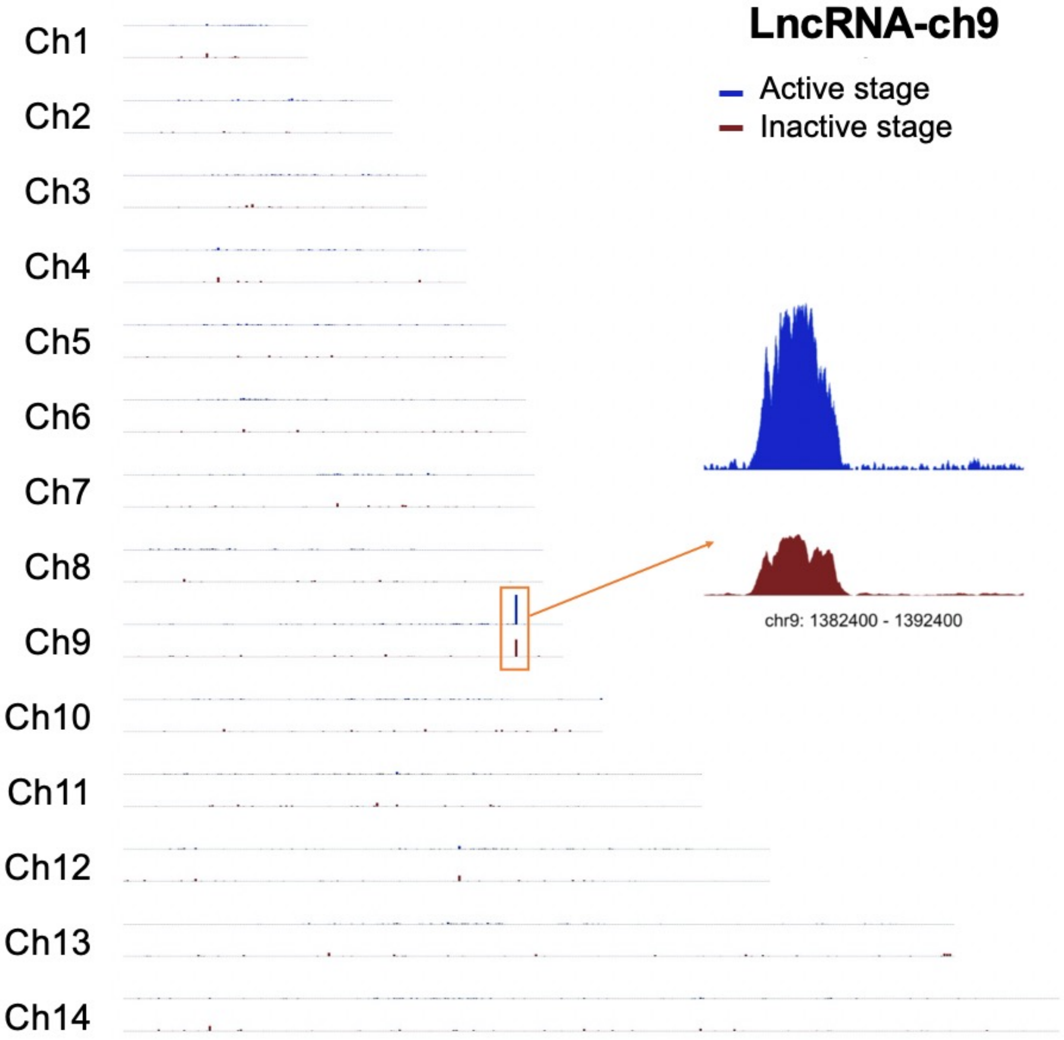

Supplementary Fig. S5

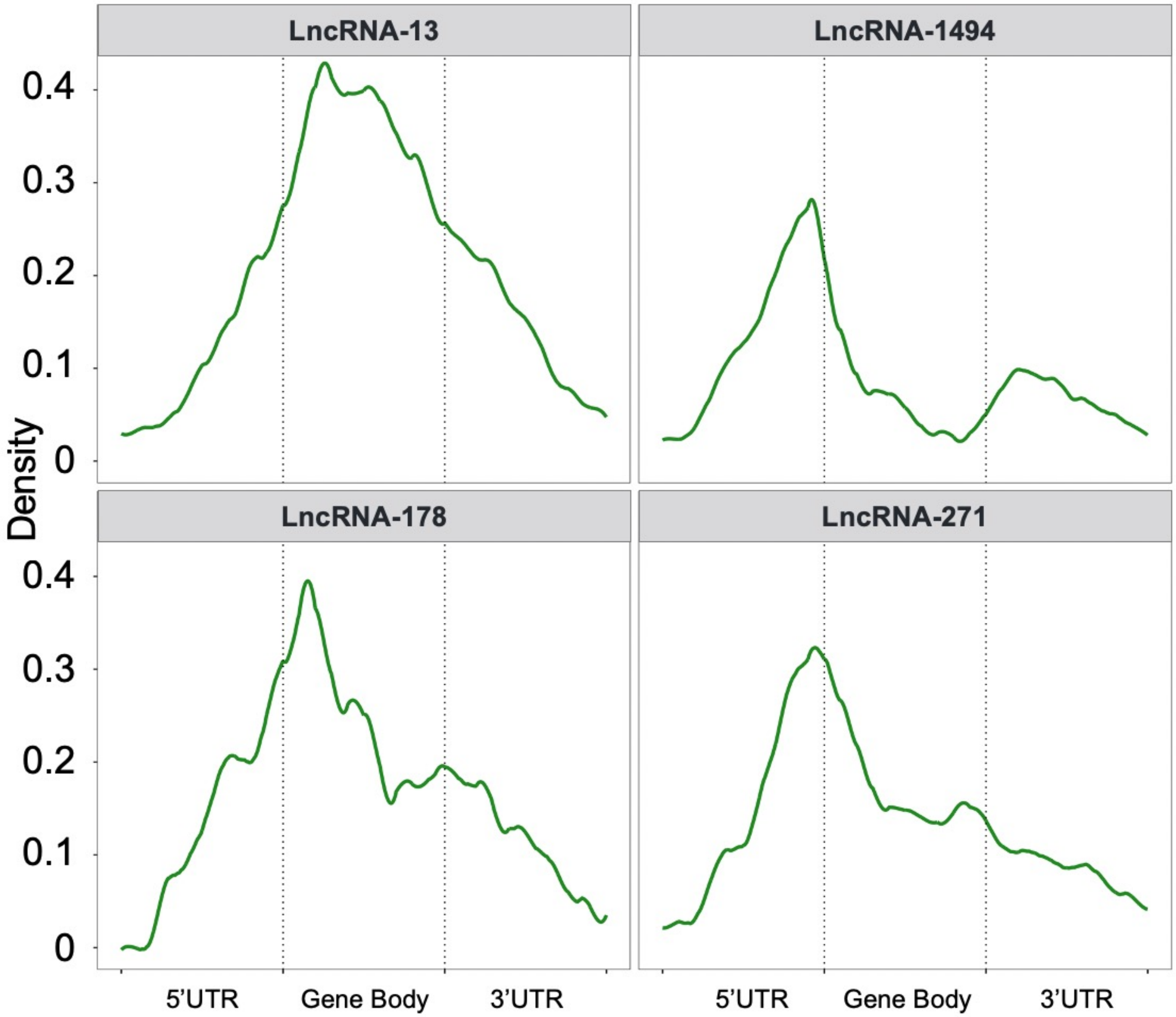

Supplementary Fig. S6

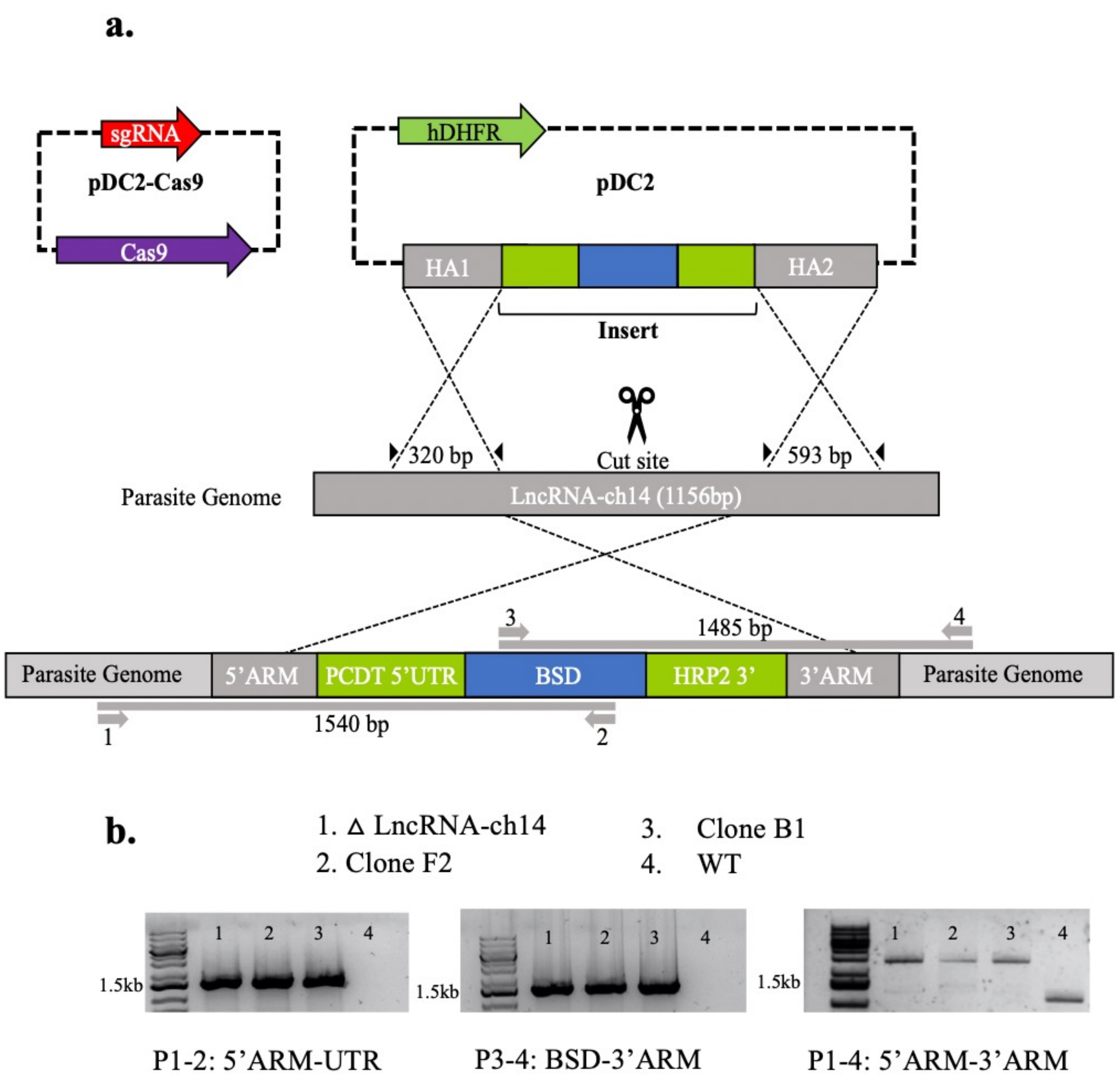

Supplementary Fig. S7

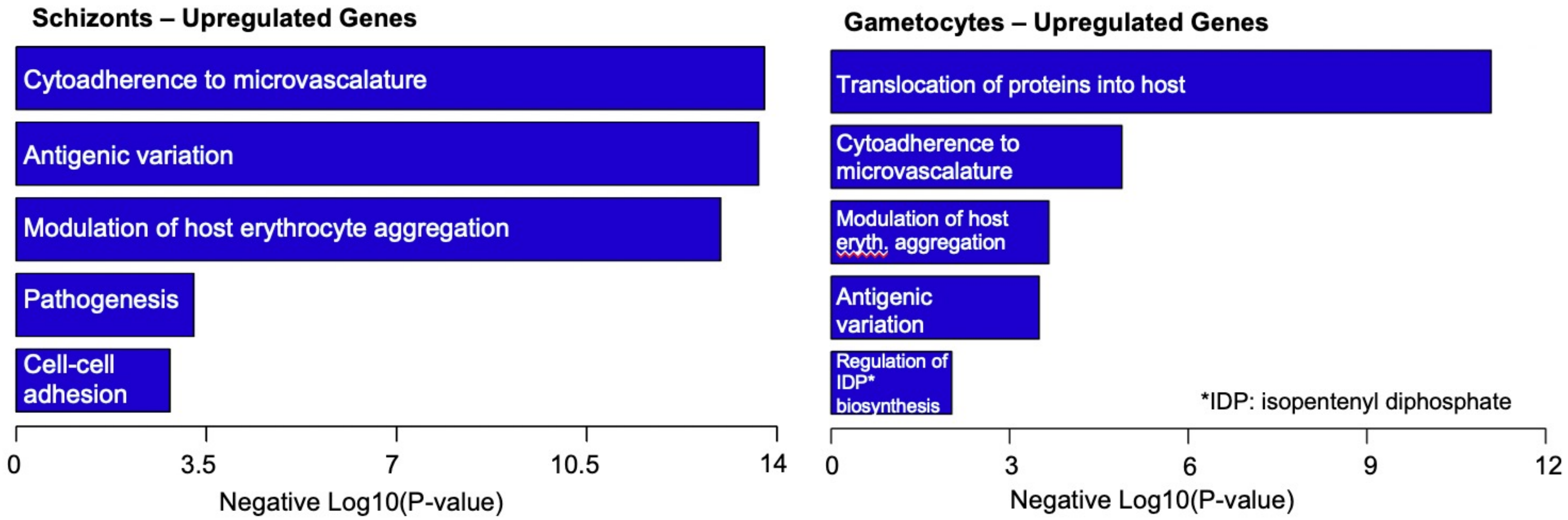
